# The molecular arsenal of the key coastal bioturbator *Hediste diversicolor* faced with changing oceans

**DOI:** 10.1101/2024.09.20.614147

**Authors:** Kaylee Beine, Lauric Feugere, Nichola Fletcher, Megan L. Power, Liam J. Connell, Adam Bates, Jiao Li, Michael R. Winter, Graham S. Sellers, Luana Fiorella Mincarelli, Sofia Vámos, Jennifer James, Hannah Ohnstad, Helga Bartels-Hardege, Daniel Parsons, Trystan Sanders, Ruth Parker, Stefan G. Bolam, Clement Garcia, Martin Solan, Jörg Hardege, Jasmin A. Godbold, Katharina C. Wollenberg Valero

## Abstract

The importance of infaunal bioturbators for the functioning of marine ecosystems cannot be overstated. Inhabitants of estuarine and coastal habitats are expected to show resilience to fluctuations in seawater temperature and pH, which adds complexity to our understanding of the effects of global change drivers. Further, stress responses may be propagated through chemical cues within and across species, which may amplify the costs of life and alter species interactions. Research into the molecular mechanisms underlying this resilience has been limited by a lack of annotated genomes and associated molecular tools. In this study, we present the first chromosome-level, annotated draft genome of the marine ragworm *Hediste diversicolor*, specifically mapping genes important for chemical communication, sensing and pH homeostasis. Using these resources, we then evaluate the transcriptomic and behavioural responses of two distinct populations — one field-sampled from Portugal (Ria Formosa) and one laboratory-acclimated and -bred from the United Kingdom (Humber) — to changes in seawater pH, temperature, and odour cues from a low pH-stressed predator. Both populations displayed adaptive responses to future oceanic conditions, with targeted acid-base regulation in the Ria Formosa population experiment, and broader changes in metabolism and growth genes in the Humber population experiment. Chemical cues from stressed fish predators induced genes related to Schreckstoff biosynthesis in ragworms. Additionally, under future ocean conditions including increased temperature, the Humber population exhibited signs of cellular stress and damage. Our findings using the new annotated genome offer novel insights into the molecular arsenal of acid-base regulation which aids in predicting the impacts of an increasingly acidified and unstable ocean, and to transfer this knowledge to investigate these mechanisms in species with less tolerance.

## Introduction

Coastal areas represent diverse habitats, with often high productivity and biomass production (Henseler *et al*., 2019), and are considered to be of high importance for “blue carbon” sequestration (Mcleod *et al*., 2011). Their heterogeneity also means that organismal communities inhabiting these ecosystems face major physiological challenges due to fluctuations in environmental conditions such as temperature, pH, salinity, or water levels through tides (Gunderson, Armstrong and Stillman, 2016). As an extension of the temperature variability hypothesis, marine life inhabiting these unstable habitats could be expected to be adapted to fluctuating temperatures (Dobzhansky, 1950; Stevens, 1989), including pH fluctuations (Thompson, Crowe and Hawkins, 2002; Hofmann *et al*., 2010). Such adaptations would convey resilience to current and, potentially, to future environmental challenges (Vargas *et al*., 2022). Yet, despite the rapidly intensifying frequency of extreme events due to anthropogenic climate change (IPCC Core Writing Team, 2023), many of the potential responses of marine infauna to environmental fluctuations remain understudied. This especially concerns the molecular mechanisms governing physiological adaptations to environmental fluctuations (Hofmann and Todgham, 2010; Somero, 2010, 2012; Lee *et al*., 2020; Strader, Wong and Hofmann, 2020; Thomas, Ramkumar and Shanmugam, 2022). This is additionally complicated by the fact that reference genomes, and consequently molecular tools to investigate these responses with, are not yet available for the majority of species.

As a consequence of rising anthropogenic carbon dioxide emissions (IPCC Core Writing Team, 2023), the oceanic pH is steadily dropping (Doney *et al*., 2009), and both seasonal (Landschützer *et al*., 2018) and short-term ocean acidification extreme (OAX) events which are often coupled with marine heatwaves (MHW-OAX events) (Burger, Terhaar and Frölicher, 2022), are becoming increasingly common. Within the organismal adaptive range, pH fluctuations are generally tolerated, but deviations in magnitude or timing can have detrimental consequences (Thompson, Crowe and Hawkins, 2002; Hofmann and Todgham, 2010; Hofmann *et al*., 2010; Duarte *et al*., 2013). During early-life stages, acidified pH is known to affect development and growth (Doo *et al*., 2012; Hue *et al*., 2022), carbonate shell development and -homeostasis (Parker *et al*., 2013), animal physiology (Melzner *et al*., 2020), fecundity (Kurihara, Shimode and Shirayama, 2004), redox homeostasis (Shi and Li, 2024), the immune system (Shi and Li, 2024), bioavailability of algal toxins (Roggatz *et al*., 2019), and metabolic and cardiorespiratory rates (Heuer and Grosell, 2014). It can also affect behaviour (Godbold and Solan, 2013; Dodd *et al*., 2015; Sanders, Solan and Godbold, 2024), with the caveat that an overall effect size is difficult to quantify (Clements *et al*., 2022; Esbaugh, 2023), possibly due to differences between study methods and ecological circumstances (Hardege *et al*., 2024). In addition, organismal interactions through chemical communication can be altered through acidified pH at different stages, from sender to signalling compound to receiver (Roggatz *et al*., 2022). For example, we have recently shown that changes in abiotic parameters in senders induces stress responses in receivers of chemical cues, including heat-stress related communication and stress propagation in developing zebrafish (Feugere, Scott, *et al*., 2021; Feugere *et al*., 2023), and pH-drop induced stress responses in marine ragworms *Hediste diversicolor* induced by both low pH-stressed conspecifics and sea bream, *Sparus aurata* (Feugere, Angell, *et al*., 2021).

Despite the wealth of studies on ocean acidification, data on the molecular basis of acid-base regulation during fluctuating conditions and abnormal exposure to pressures such as climate-change induced upwelling events, SST associated pH changes, or ocean acidification in general, are still scarce across most species. In marine ectotherms, increasing environmental *CO*_2_ leads to a rise in body fluid *CO*_2_ and an acidosis (drop in pH) of extra- and intracellular fluids (Melzner *et al*., 2020). In strong acid-base regulators (e.g. most polychaetes), this acidosis is rapidly buffered by accumulating bicarbonate ions (HCO_3_^-^), obtained either from the hydrolysis of CO_2_ catalysed by the enzyme carbonic anhydrase *(Wäge et al., 2016)*, metabolic decarboxylation of amino acids (Langenbuch and Pörtner, 2002) and/or branchial bicarbonate uptake utilising HCO₃⁻-Cl⁻ anion exchangers (Heuer and Grosell, 2014). Excess protons also require excretion via respiration and/or renal systems through cation exchangers, or energy-demanding ATPases such as V-type H^+^-ATPases (Alberts *et al*., 2002; Tresguerres, 2016). This increased energy requirement for ion and acid-base regulation may result in the above-mentioned negative consequences or trade-offs with other physiological processes (Widdicombe and Spicer, 2008).

Marine ragworms (class Polychaeta) constitute a major component of the coastal sediment infauna, for example, they have been reported to have an annual mean biomass of 66.244 g/m^2^ in a Portuguese estuary (Carvalho *et al*., 2013). Ragworms have key ecological roles in coastal ecosystem functioning, within coastal food webs (Masero *et al*., 1999), and by oxygenating the sediment through bioturbation and organic matter breakdown, both of which are crucial for maintaining benthic ecosystem health (Banta *et al*., 1999; Mermillod-Blondin, 2011). Absence of bioturbators such as ragworms has been associated with anoxia in sediment through collapse of sediment mixing, which was a characteristic of the end-Permian mass extinction (Hofmann *et al*., 2015). Two polychaete species, *Platynereis dumerilii* and *Capitella teleta,* have become laboratory model organisms for developmental and neurobiological studies with available reference genomes (NCBI:txid283909 for *C. teleta* and NCBI:txid6359 for *P. dumerilii* (Mutemi *et al*., 2024) and a wealth of associated molecular tools (Seaver, 2016; Özpolat *et al*., 2021), whereas research in non-model ragworms such as *Hediste diversicolor* (NCBI:txid126592) often focuses on ecology (García-Arberas and Rallo, 2002; Gilbert *et al*., 2021), feeding and diet (Aberson, Bolam and Hughes, 2016), and ecotoxicology (Fonseca *et al*., 2017; Ghribi *et al*., 2019; Silva, Pires, *et al*., 2020).

*Hediste diversicolor,* formerly *Nereis diversicolor,* is a common polychaete ragworm, inhabiting shallow marine and brackish waters of European and North American coastlines (Scaps, 2002; Catalano *et al*., 2012). They serve as biological models in ecotoxicological research due to their high sensitivity and fast reaction to contaminants (Mouneyrac, Perrein-Ettajani and Amiard-Triquet, 2010; Silva, Pires, *et al*., 2020). These studies have included pharmaceuticals (Nunes *et al*., 2016), metals (Freitas *et al*., 2017; Silva, Pires, *et al*., 2020), micro- and nanoplastics (Gomiero *et al*., 2018; Silva, Oliveira, *et al*., 2020), polyaromatic hydrocarbons (Catalano *et al*., 2012) and seawater acidification (Freitas *et al*., 2016, 2017).

Acidification of seawater pH has been shown to affect metabolic processes in *H. diversicolor,* such as higher carbonic anhydrase activity, lower energy reserves and higher metabolic rate (Freitas *et al*., 2016). However, most current studies using *H. diversicolor* use either candidate enzyme or metabolic product approaches (Freitas *et al*., 2016, 2017; Nunes *et al*., 2016; Sokołowski, Brulińska and Sokołowska, 2020; Daniel *et al*., 2022) rather than an -omics focus. The molecular mechanisms underlying pH homeostasis and short-term regulation in marine ragworms therefore remain largely unexplored, yet understanding these processes is necessary for predicting their responses to future ocean changes.

Here, we provide a new annotated chromosome-level resolution draft genome for *H. diversicolor* based on PacBio long read sequencing combined with Omni-C chromosome conformation capture. We use this new reference genome to elucidate 1) the phylogenetic position of *H. diversicolor* among genome-sequenced marine polychaetes, 2) the characterization of environmental sensing and ion and acid-base regulation genes within the genome, and 3) to provide exemplary baseline data on the behavioural and transcriptomic responses of two geographically distinct populations of *H. diversicolor*, one from the UK (Humber Estuary (“Humber” hereafter) and the other from Portugal (Ria Formosa), to short-term pH changes either alone, combined with higher temperature, or signalled through chemical cues by low pH-exposed sea bream (*Sparus aurata*). Given its ecological role as a bioturbator inhabiting variable environments, we expected *H. diversicolor* to be able to mount an adaptive response to these fluctuations.

## Materials and Methods

### Genome sequencing and annotation

*Hediste diversicolor* were collected from the Humber estuary (53.7144° N, 0.4458° W) and subsequently inbred within a small group in the laboratory at the University of Hull over two generations in control conditions. In February 2021, four adult F2 ragworms (example Figure 1A) were flash-frozen in liquid nitrogen and sequenced at the Cantata Bio (formerly Dovetail Genomics, California) sequencing facility with Omni-C and PacBio methods and a combined assembly was generated using *bwa* (https://github.com/lh3/bwa). For annotation, we ran the BRAKER2 pipeline for gene prediction (Brůna *et al*., 2021) against both a concatenated set of nine transcriptomes obtained from the Ria Formosa *H. diversicolor* population and on protein evidence from Polychaeta in UniProt - OrthoDB (Kuznetsov *et al*., 2023; UniProt Consortium, 2023) followed by applying a custom script to collapse unlikely gene models. Additional annotations resulted from scans for noncoding RNAs with *Infernal* (Nawrocki and Eddy, 2013; Kalvari *et al*., 2021) and the single-to multi-exon gene ratio (SEG:MEG ratio, also referred to as “mono-exonic/multi-exonic gene counts” or mono:multi ratios (Vuruputoor *et al*., 2023) was calculated to serve as a quality check of the annotation and was determined across all protein-coding genes (for details, see Supplementary Methods).

### Synteny and phylogenetic analysis

Whole genomes for polychaetes were downloaded from GenBank (*Platynereis dumerilii* GCA_026936325.1*, Alitta virens* GCA_932294295.1*, Capitella teleta* GCA_000328365.1*, Nereis procera* GCA_032360585.1). Since all these marine polychaetes were in different genera, whole-genome alignment and synteny analysis was attempted but not successful. Instead, BUSCO (v 5.6.1) (Simão *et al*., 2015; Manni *et al*., 2021) was used to find the universal single copy orthologs between the species including our own *H. diversicolor* assembly, using the Eukaryota database. Single-copy orthologs were used to determine synteny between the species. *Capitella teleta* was not similar enough to establish synteny with the other species. *Nereis procera* had a BUSCO score of 3.5% and only 8 genes were considered to be single-copy genes, therefore synteny could not be established either. Synteny for *H. diversicolor*, *P. dumerilii*, and *A. virens* based on the chromosomal locations of BUSCO genes were analysed and plotted using R packages *GeneSpace* (Lovell *et al*., 2022) and *circlize (Gu et al., 2014)*. In *GeneSpace,* an orthogroup is defined as a group of genes that are orthologous across different species. Orthogroups are identified using OrthoFinder, a tool that groups genes from multiple species into orthologous groups based on sequence similarities and phylogenetic relationships. Bayesian phylogenetic inference (BI) was performed on the 299,373 nucleotides from the BUSCO gene alignment with BEAST v1.8.4 (Suchard *et al*., 2018).

### Acid-base regulation and other sensory genes in the genomes of H. diversicolor

Due to the influence of acidified seawater pH on functions like acid-base regulation, behaviour and chemical communication, we were interested in specifically annotating genes with known function in neuronal signalling, environmental sensing, and pH sensing and -homeostasis. For this purpose, we searched the Gene Ontology Biological Process database (Ashburner *et al*., 2000; Gene Ontology Consortium *et al*., 2023) for genes with related terms resulting in: GO0004089 Carbonate dehydratase activity (equivalent to carbonic anhydrase); GO0030641 Regulation of intracellular pH; GO0006885 Regulation of pH; and GO0007635 Chemosensory behaviour (for complete list of terms and genes see Electronic Supplementary Datasheet tab 6). The search taxonomy was reduced to *Caenorhabditis briggsae* and *C. elegans* as closest relatives to marine ragworms in this database and matching gene sequences were obtained from NCBI or UniProt. Additional signalling genes were obtained from two publications, one on neuropeptide signalling via G-protein coupled receptors (GPCRs) in *Platynereis dumerilii* (Bauknecht and Jékely, 2015); accession numbers GenBank:KP293941–KP294026, and another on peptidergic brain signalling in larval *P. dumerilii*; accession numbers GenBank:KP420212–KP420214), (Williams *et al*., 2017). Input *fasta* files were generated from these gene sequences and searched against the *H. diversicolor* draft genome via *discontinuous megablast* algorithm using BLAST+ (Camacho *et al*., 2009). Gene models matching these gene entries in *H. diversicolor* were annotated in a separate field in the genome annotation GTF file “*sensory_function*”. In addition, the GTF file was scanned for already annotated instances of the known receptor gene classes ABC transporters, GABAα receptors, NMDA receptors, OTOF, OTOP, and OTOG (mechanoreception), SC5A8 (lipid transport), TAAR receptors (amino acid sensing), TRPV receptors (environmental sensing), and XKR scramblases (possible social signalling (Feugere *et al*., 2023) and, where found, these were likewise annotated in the *sensory_function* column. A karyotype map of *H. diversicolor* with annotated sensory and signalling genes was drawn with *ChromoMap* in R (Anand and Rodriguez Lopez, 2022), and the most common genes were counted.

### Burrowing behaviour of H. diversicolor from the Humber in response to future ocean conditions

Specimens of *H. diversicolor* were collected from the Humber Estuary, United Kingdom (53.7144° N, 0.4458°W) between December 2020 and April 2021 and transported to the University of Hull within 2 hours of collection. Animals were placed into two separate banks. Each bank was connected to a continually filtered, closed loop system consisting of 10cm depth fine sand sediment and 10 cm circulating water (salinity 22 ppt) and subdivided into five tanks to allow for independent replicates. Each system was continually aerated and ragworms fed twice weekly on crushed Tetra TabiMin. Light conditions were maintained at 12L:12D with a moon cycle tracking the natural cycle. Temperature and pH in both banks were manipulated to dynamically mimic both current day oceanic conditions (**current**) and forecast future ocean conditions (**future** (IPCC Core Writing Team, 2023). The pH was controlled via the Loligo CapCTRL system through addition of CO_2_ gas. Temperatures were controlled using a DC300 aquarium chiller, since all experimental temperatures were below room temperatures. Each aquarium was programmed to follow a fluctuating pH in line with current of predicted future diurnal cycles (Pacella *et al*., 2018). **Current** day conditions were: average pH 8.2; (as per the National Bureau of Standards scale) temperature ranging from 13°C in winter to 16°C in the summer. **Future** conditions were: average pH 7.7 with a temperature of +3°C over those of the current condition tanks. Supplementary Figure 1 shows a graph of natural pH fluctuation of the Humber estuary near Grimsby in Spring and Summer of 2021 with an average spring pH of 7.57 and an average summer pH of 7.95. Supplementary Figure 2 shows an example for measured diel pH fluctuation in these two long-term treatments. Animals from both conditions were then used for behavioural bioassays and RNA sequencing as detailed below. For behavioural assays of the Humber population, in October 2021, animals were collected from each condition and put in individual glass beakers (6 cm diameter) containing 1 cm depth of the same sediment and water from the corresponding aquaria. Ragworms were timed from being added to the beaker to burial of the complete head segment, for a maximum of 300 seconds (**current** n = 30; 51 observations; **future** n = 30, 58 observations). Within minutes, a subset of ragworms were then used for a second and third behaviour test consecutively, with ragworm use # recorded as a covariate to see how subsequent handling affected this burrowing behaviour and to account for pseudoreplication. All animals were returned to their original tanks after experiments were completed. To additionally test for burrowing behaviour on instant exposure to short-term altered pH, a further n=39 ragworms with 53 observations acclimated to the current treatment were transferred into water from the treatment (**current → future**) and their burrowing behaviour was timed as above. This was likewise done with using animals acclimated to future conditions treatment tested in current conditions water n = 45 ragworms with 73 observations (**future → current**).

### Burrowing behaviour of H. diversicolor from Ria Formosa in response to low pH

The second population of *H. diversicolor* was obtained from a local supplier (Valbaits) in Ria Formosa, Portugal during October 2019 and 2023 with the pH of their natural habitat ranging from 7.75 to 8.0 (Nascimento *et al*., 2021). Animals underwent acclimation for three to four days at a pH of ∼8.1 (as per the National Bureau of Standards scale) in large, flow-through open-circuit communal tanks located outdoors at the Ramalhete Marine Station (CCMAR, Faro, Portugal; continuous flow of 0.05 L/min). Animals were not fed during this period. The pH levels were maintained using a direct CO_2_-controlled system which continually monitors and adjusts the pH (for details see (Sordo *et al*., 2016, 2018; Gregório *et al*., 2019)). Two seawater conditions were used from two header tanks, as described in (Feugere, Angell, *et al*., 2021): a control pH (“**current**”, pH = 8.136) and acidified (“**future pH**”, pH = 7.768). Temperature and light regimes followed the ambient profile in Faro. Burrowing assays were performed as outlined above for the Humber population, except that the dishes were 9 cm in diameter. Methods are described in detail in Feugere et al. (Feugere, Angell, *et al*., 2021). Behavioural tests were performed for these ragworms acclimated in **current** (n = 24, with 51 observations) and **future pH** (n = 51, with 78 observations) conditions. In addition, the response of ragworms was tested in the **current → future pH** (n=9, with 36 observations) and **future pH → current** (n=24, with 24 observations) condition. Here, the number of worm reuses was likewise considered as a predictor variable.

### Burrowing behaviour data analysis

In the case that ragworms burrowed the head within 300 s, this was dubbed a “success”. Data for each population was analysed separately due to the differences in experimental setups, and specifically, in acclimation conditions for the Humber population (Supplementary Table 1). “Time-to-success” was modelled for the time to burrow the head in *H. diversicolor* depending on the four experimental conditions using Cox proportional hazard (Coxph) models implemented in the *survival* R package v3.5-7 (Therneau and Grambsch, 2000; Therneau and Lumley, 2015), defining the event as successfully burrowing the head. Animals not burrowing their heads within 300 s were censored. The exponentiated estimates (hazard ratios) from the Coxph models were expressed as “success ratio” (Supplementary Table 2) and visualised using the *sjPlot* R package v2.8.15 (Lüdecke, 2018). Probability of successfully burrowing the head over time was visualised with Kaplan-Meier curves using the *survminer* R package v0.4.9 (Kassambara et al. 2021) and burrowing speed represented with box plots and tested for differences with pairwise Wilcoxon tests (Supplementary Table 3). The addition or not of covariates (worm reuse, source temperature, test temperature, and the difference between source and target temperature) was determined by using the Coxph model with the lowest second-order (small-sample size) Akaike Information Criterion (AICc) based on the R package *AICcmodavg* v2.3-3 (Mazerolle, 2023).

### Transcriptomics of H. diversicolor

Adult field-sampled ragworms were acclimatised in 5 L glass tanks with natural sand alimented by control system water at pH = 8.088, temperature = 21.9 °C, salinity = 35.7 psu, and O_2_ = 7.3 mg. L^-1^ (note the small difference to the behaviour experiment) for approx. five hours before the commencement of the experiment. The experiment was conducted in October 2019 at ambient temperature (∼22.5 °C) and under natural light conditions. Experimental treatments aimed to test the direct exposure of ragworms to lowered pH (**current → future pH**), but also the exposure to chemical cues (“Stressed Fish Metabolites SFM”) from fish that had additionally been exposed to lowered pH (**current → SFM**) (see (Feugere, Angell, *et al*., 2021) for method details). The three conditions were: 1) control pH = 8.1 (pH = 8.075); 2) low pH 7.6 (pH = 7.623); and 3) water conditioned by a low pH-exposed fish. Conditioned fish water was obtained by placing one gilthead sea bream *Sparus aurata* (weight: 820 g) at low pH = 7.6 for 30 minutes in a 15 L bucket with an air pump. Water conditioned with the gilthead sea bream was provided by the Ramalhete marine station. 50 mL of this conditioned fish water at pH 7.6 was diluted in 450 mL control system water at pH 8.1, with a resulting pH of 8.02 (for details, see Feugere et al., 2021). Just before the beginning of the 1-hour exposure, a volume of seawater necessary to expose 5 ragworms was prepared per condition in a 500 mL container. Salinity was measured, directly in the stock solution of conditioned seawater and just before the beginning of the exposure, as 37 ppt in all three conditions, and as 22.6 ℃ in the SFM condition and 22.5 ℃ in the control and low pH conditions. For each condition, five ragworms were weighted in seawater (weight = 0.78 ± 0.37 g) and were individually placed in 50 mL centrifuge tubes containing approx. 25 g of clean sand (10 mL) and completed with approx. 40 mL of medium (tube filled to 50 mL) of each experimental treatment (for details, see Supplementary Methods). 65 µL RNA samples meeting at least the minimum concentration and quality requirements were processed by Edinburgh Genomics who generated cDNA libraries using the TruSeq stranded mRNA kit and sequenced reads using a Illumina NovaSeq 6000 50PE sequencer.

Ragworms from the Humber population were much smaller than those from Ria Formosa. Therefore, it was not feasible to obtain enough RNA from the head and it was decided to use whole-body specimens instead. Directly upon a one-hour exposure in **current** (used as the control), **current → future pH**, and **current → future**, n = 5 ragworms per each of these three treatments were flash-frozen and stored at -80°C in RNAlater^®^. RNA was extracted as described in Supplementary Methods and sequencing was done by Liverpool Genomics (UK) (for details, see Supplementary Methods).

### Transcriptomic data analysis

Raw RNA reads were processed to remove adaptors and low quality reads using *fastp* v0.23.4 (Chen *et al*., 2018), with deduplication disallowed. *FastQC* v0.11.9 (https://www.bioinformatics.babraham.ac.uk/projects/fastqc/) was used to assess read quality before and after trimming sequences. Paired-end reads were then mapped to the *H. diversicolor* genomes using *HISAT2* v2.0.0 (Kim *et al*., 2019) and results were sorted and converted to BAM format using *SAMtools* v1.10 (Li *et al*., 2009). Reads were quantified with *Stringtie* v2.1.6 (Pertea *et al*., 2015) and read count matrices generated from the Stringtie output using the prepDE.py script (https://github.com/gpertea/stringtie/blob/master/prepDE.py). Differential gene expression analysis was performed using *DESeq2* v1.44.0 (Love, Anders and Huber, 2013) in R v4.4.1 using *Bioconductor* v3.19 (Morgan, 2024). For the Ria Formosa population, gene expression data were processed with treatments and weight in the design matrix, and pairwise comparisons were performed **current → future pH** and **current → SFM** with current condition samples as controls. For the Humber population, given differences in library across samples, the counts were normalised using the Trimmed Mean of M-values (TMM) method implemented in the *edgeR* package (Robinson, McCarthy and Smyth, 2010). The TMM normalisation adjusts for library size variations by calculating normalisation factors, which were subsequently applied to the raw counts. Normalised counts were converted into counts per million (CPM) to facilitate downstream analyses. Pairwise comparisons for the Humber population were **current → future pH** and **current → future.** Differentially expressed genes (DEGs) for all comparisons were visualised as volcano plots drawn using *EnhancedVolcano* v1.22.0 (Blighe, 2024). Orthologs for DEGs which were identified in BLAST searches were determined for Human, with orthoDB (Kuznetsov *et al*., 2023) and UniProt and ClueGO (Bindea *et al*., 2009) was then used to conduct GO term analysis, using median network specificity and the human genome as background. Bonferroni correction for term p-values and GO term merging to reduce GO term hierarchy. Gene Ontology databases included Biological Process, KEGG pathway, and Reactome pathway.

## Results

### The genome of Hediste diversicolor

Taxonomic clean-up of the PacBio assembly using *blobtools* (Laetsch and Blaxter, 2017) revealed that the best taxon matches of the reads were Annelida and Mollusca (Figure 1C). Most PacBio reads had a coverage of 100x, which was much higher than the envisioned 30x coverage (Figure 1C). The link density histogram of read-to mate-pairs in Omni-C scaffold assembly in Figure 1B shows 14 major TADs (topologically associated domains of chromatin), which represents possible chromosomes in the haploid genome (2n=28). Table 1 shows the statistics of the combined PacBio and Omni-C genome assembly for *Hediste diversicolor*. The haploid genome size is estimated as 0.95 Gb, which is similar to that of other marine polychaetes (*Platynereis dumerilii* -1.47 Gb, (Mutemi *et al*., 2024), 0.32 Gb *Capitella teleta* (Simakov *et al*., 2013); 0.61 Gb *Alitta virens* (Fletcher *et al*., 2023). BUSCO found 92.10% of essential genes, with 13 reported missing. The other polychaete species analysed were *C. teleta* (95.70% essential genes, 3 missing); *A. virens* (96.90% essential genes, 2 missing), *P. dumerilii* (96.40% essential genes, 4 missing), and *Nereis procera* (3.5% essential genes, 137 missing, in following excluded from synteny analysis). The total Transposable Element (TE) content amounted to 52.98% of the genome with 26.27% being of unknown classification (Table 1). The final genome annotation contained 27,839 gene models of which 14,840 did have BLAST hits and 12,999 did not (Figure 1E). To this final version of the annotation, we appended 5338 instances of noncoding RNA genes and 25 miRNAs predicted with *Infernal* (Table 1; Electronic Supplementary Datasheet tab 1, 2 and 3). The number of CDS per gene model was positively correlated with cumulative CDS length showing that longer genes also consisted of more exons. Notably, non-BLAST-annotated genes had fewer exons than annotated genes (Figure 1D). The SEG:MEG ratio was 15.6% SEGs, with the majority of non-BLAST-annotated genes being relatively short with a low number of exons. Both *H. diversicolor* and *A. virens* genomes supported the presence of n=14 chromosome-scale scaffolds, while both the *H. diversicolor* and the *P. dumerilii* genomes had additional smaller scaffolds, possibly representing genomic regions with uncertain chromosomal association (Mutemi *et al*., 2024).

**Figure 1.**
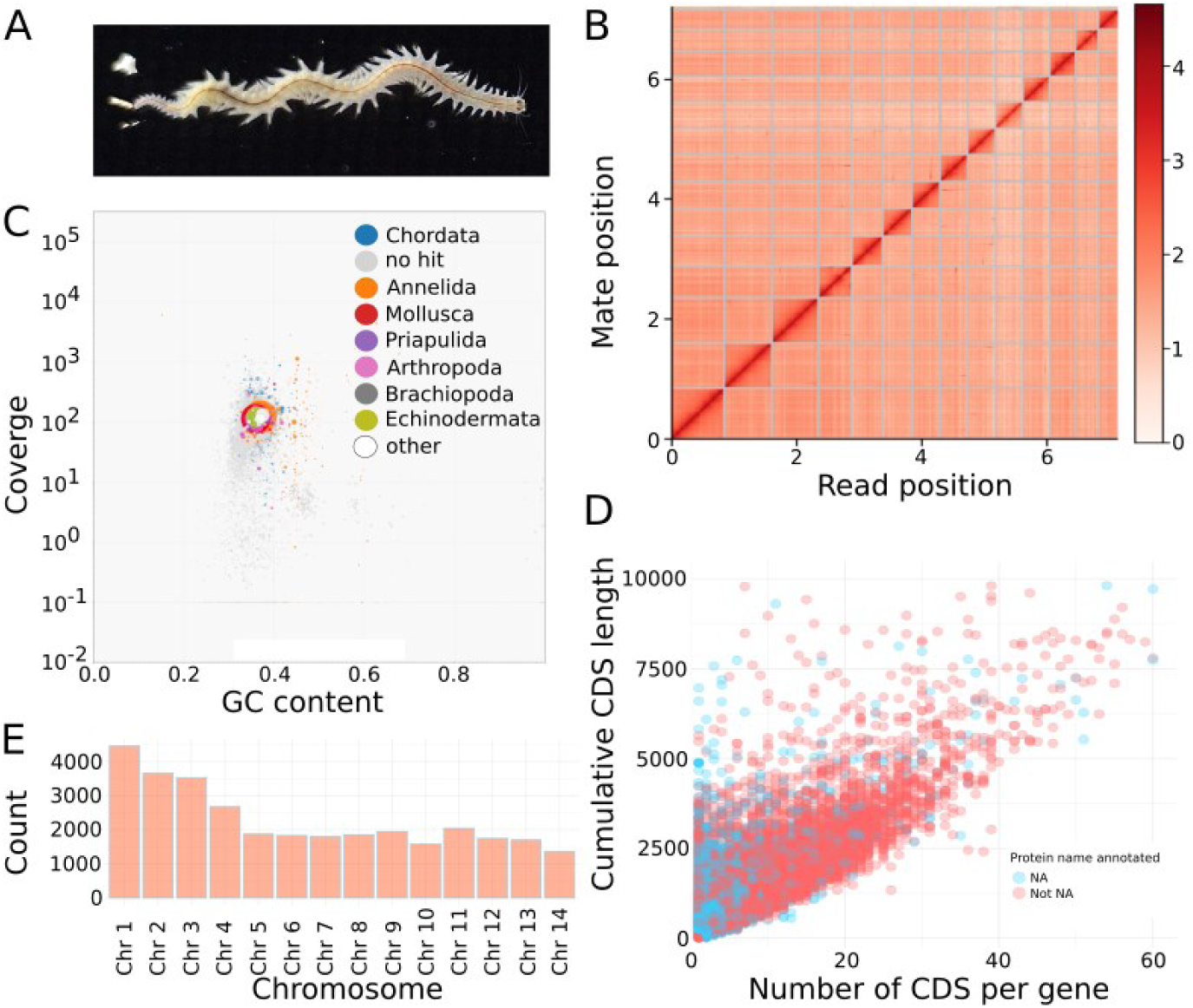
Clockwise: A) example illustration of *Hediste diversicolor* sequenced for chromosome conformation capture. B) Link density histogram of read-to mate-pairs in Omni-C scaffold assembly showing 14 chromosome-level assemblies in haploid genome (n=14; 2n=28). Dark red squares on the diagonal represent TADs (topologically associated domains of chromatin). C) Blob Plot of PacBio sequencing assembly with taxon matches in colours, with best matches to Annelida and Mollusca. Most PacBio reads had a coverage of 100x. D) Correlation plot between number of coding sequences in gene models and the cumulative length of the CDS of the gene model (axes truncated). Gene models represented by blue dots have no BLASTp annotation and red coloured gene models have BLASTp annotation. It is evident that non-annotated gene models (and therefore likely also some false positives), are concentrated in shorter genes with fewer CDS. Non-annotated Single-exon genes with a CDS length less than 250bp were excluded from the annotation. E) Histogram of number of genes per chromosome-level scaffold for the 14 major scaffolds.

**Table 1.** Summary of the Genome Assembly and Annotation for *Hediste diversicolor* with statistics performed on the High-Rise genome assembly combining PacBio and Omni-C technologies. SEG:Single-exon genes; MEG: multi-exon genes. TAD - Topologically associated domain of chromatin (Electronic Supplementary Datasheet tab 1 and 2).

|  |  |  |
| --- | --- | --- |
| **Genome assembly (Dovetail HiRise Assembly)** | Estimated haploid genome size | 951,181,829 bp = 0.95 Gb |
|  | N50 scaffold length | 44.26Mb |
|  | L50 | 9 |
|  | N90 scaffold length | 42,955Mb |
|  | L90 | 1,291 |
|  | No. of TADs (chromosome level scaffolds) | 14 |
| **BUSCO (genome)** | Complete | 235 (92.10%) |
|  | Complete and single-copy | 226 (88.63%) |
|  | Duplicate | 9 |
|  | Fragmented | 7 |
|  | Missing | 13 |
|  | Total BUSCO groups searched | 255 |
| **Protein-coding genes** | No. of gene models | 27,839 |
|  | Functionally annotated | 14,840 |
|  | Mean gene length | 6395 bp |
|  | Mean no. of exons per gene | 6 |
|  | Mean exon length | 187 bp |
|  | Mean CDS per gene length | 1124 bp |
|  | SEG:MEG percentage | 15.6% |
| **Transposable elements** | LTR-retrotransposons | 17.33% |
|  | LINEs | 8.75% |
|  | DNA-transposons | 6.35% |
|  | Rolling circles | 0.06% |
|  | Unclassified/unknown | 26.27% |
| **Classes / n loci of noncoding RNAs Predicted *ab initio*** | tRNA | 2026 |
|  | Histone 3' UTR stem-loop | 1596 |
|  | 5SrRNA | 1056 |
|  | U4 or U2 spliceosomal | 154 |
|  | 5.8S rRNA | 102 |
|  | miRNA | 25 |
|  | piRNA | 5 |
|  | other | 374 |

### Phylogenomics and the sensory arsenal of H. diversicolor

Based on the BUSCO analysis, there was evidence for extensive syntenic remodelling between all three species, mostly among the largest chromosomes, with several orthogroups being found in all three species (Figure 2). The BUSCO derived Maximum Likelihood phylogeny however resulted in an expected topology, with *A. virens* being the sister clade to *H. diversicolor*, *P. dumerilii* being the next closest related species, and *C. teleta* constituting the outgroup (All nodes with Bayesian support of 1; Figure 2).

**Figure 2.**
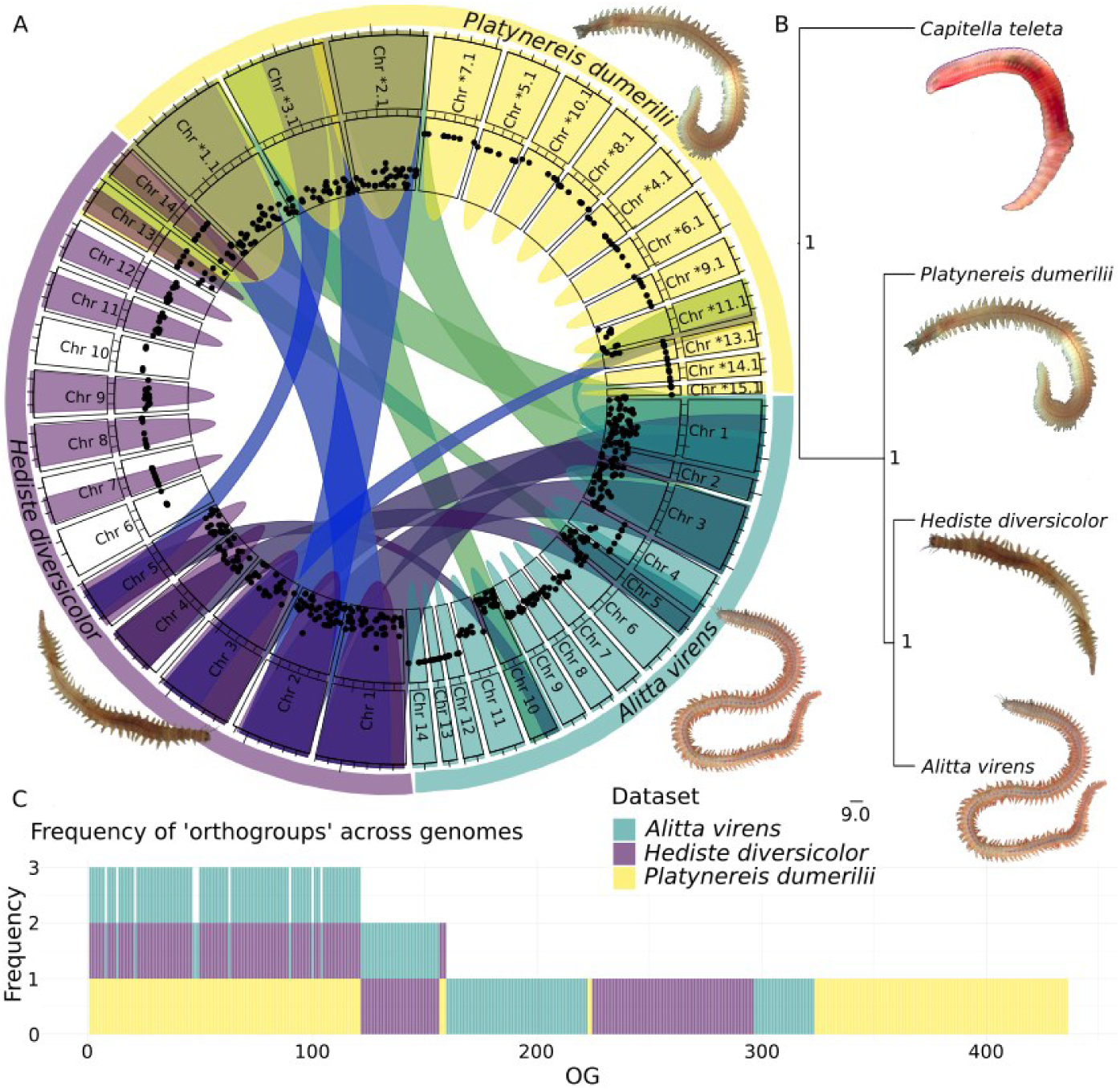
A) Syntenic regions based on BUSCO gene orthogroups (OGs) between *Alitta virens, Platynereis dumerilii,* and *Hediste diversicolor.* Data plotted by sequence of chromosome scaffold number, except in the case of *P. dumerilii,* which was based on length, abbreviation for Orthofinder identifier scaffolds set as *=JPTHN01000000. B) Phylogenetic tree created with BUSCO genes (n = 200 genes shared between the 4 species), with node support values from Bayesian inference. C) Frequency plot for which genomes each BUSCO orthogroup (sorted by ascending number) has been found in, showing that several orthogroups occur in more than two genomes, which could consequently be used for synteny pattern identification. Image copyrights: *C. teleta* Lauren Ku under CC BY-SA 4.0 licence via Wikimedia Commons (https://commons.wikimedia.org/wiki/File:Thumbnail_Capitella_teleta.jpg) and *A. virens* Yale Peabody Museum under CC 1.0 Universal licence via Wikimedia Commons. (https://commons.wikimedia.org/wiki/File:Alitta_virens_%28YPM_IZ_071022%29.jpeg). *P. dumerilii* and *H. diversicolor* are original images taken by authors.

The targeted annotation of sensory and signalling genes resulted in the identification of 240 gene annotations across the genome of *H. diversicolor* (Figure 3). Most were concentrated on the seven largest chromosomes, with chromosomes 8-10 containing the lowest number. The most frequently annotated genes were OTOP, GABR1, FMAR, GPR34 (orthoDB:810985at33208, 2325at6231, 5404226at2759, 3875139at2759) as well as other TRP channels and GPCRs (TRPV5, TRPA1, GPR29, TRPV6, GPR47; orthoDB:1003028at2759, 2910129at2759, 2325at6231, 1003028at2759, 2325at6231) as well as GABR2 (orthoDB:2970339at2759). This means that the most frequently annotated functions were mechanoreception, GABAα reception, G-protein coupled receptor, and TRP channels (Figure 3). The third most frequently annotated gene was FMAR (FMRFamide receptor; orthoDB:5404226at2759).

**Figure 3.**
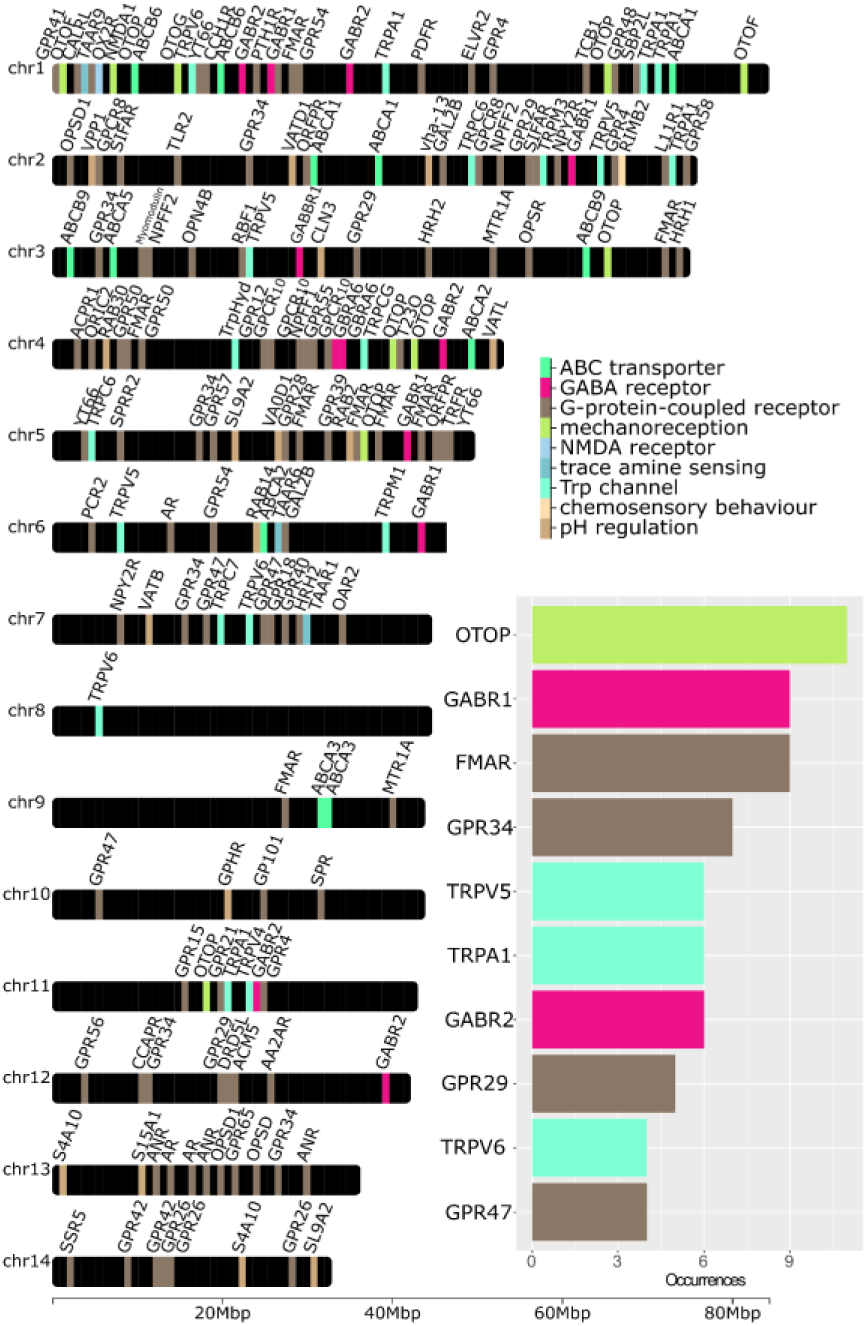
240 sensory gene loci annotated on the 14 chromosome-level scaffolds of *Hediste diversicolor*, highlighting different putative functions. Inset diagram shows number of occurrences of 10 most frequently annotated sensory function genes in the genome of *H. diversicolor*.

### Using the sensory arsenal in response to low pH: behaviour

The burrowing behaviour of both populations was impacted by future ocean conditions (Figure 4). Additionally, the prediction of the burrowing response was more accurate (i.e., lowest AICc) for Humber ragworms when considering ragworm reuses, as well as source and test temperatures (Electronic Supplementary Datasheet tab 8). In the Humber population, which was a field-collected population acclimated between 6 and 10 months to fluctuating indoor laboratory conditions, the control condition **current** ragworms took the longest to burrow, while the burrowing success ratios were significantly higher when ragworms were acclimated for months in **future** ocean conditions (p=0.005), and when switched back to current **future → current** condition (p<0.001; Figure 4 A,B,C, Supplementary Tables 2-3). Burrowing success probabilities were highest in the short-term swap to future conditions in **current → future** condition (p<0.001; Figure 4B), due to ragworms burrowing faster than in the **current** condition (p=0.012; Figure 4C, Supplementary Tables 2-3). In the Ria Formosa population, the influence of long-term acclimation to low pH was also evident with acclimated ragworms from both the **future pH** (p <0.001) and **future pH → current** conditions (p<0.001) having the highest burrowing success probability (Figure 4D,E,F), and being also much faster in burrowing (Figure 4, Supplementary Tables 2-3). In contrast, the swap from **current → future pH** in this population led to a significant decrease in both burrowing success (p<0.001) and speed (p < 0.001), which shows the opposite pattern from the behaviour of the Humber population for this condition (Figure 4, Supplementary Tables 2-3).

**Figure 4.**
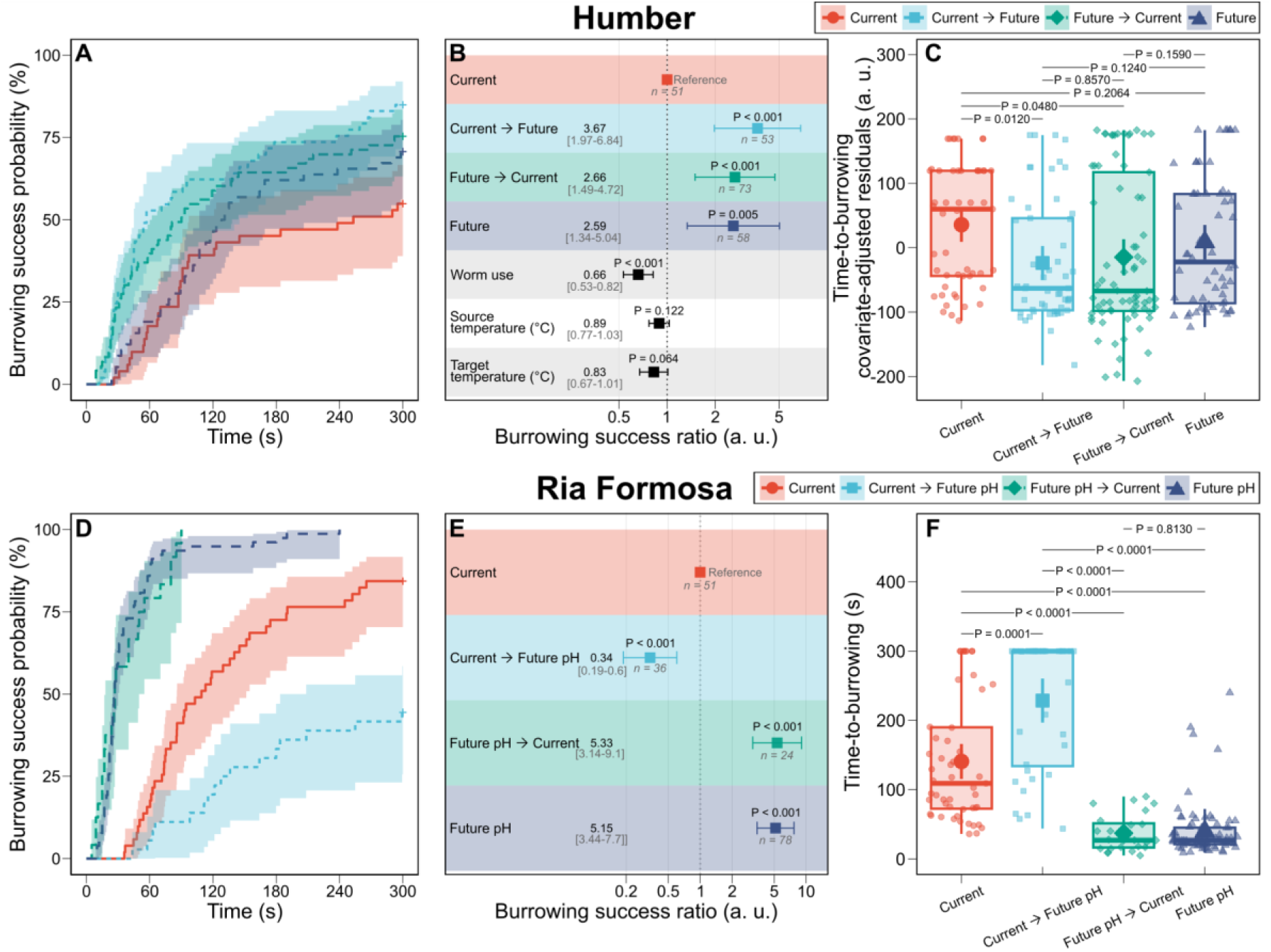
Future ocean conditions alter burrowing responses. Kaplan-Meier curves of burrowing success probability over time in *Hediste diversicolor* from the Humber (A-C, n = 235 ragworms) and Ria Formosa (D-F, n = 189 ragworms). Forest plot representing the Cox-proportional hazard models for the Humber (B; 169 events; global P < 0.0001; ΔAkaike Information Criterion [AICc] relative to null model = -24.18; concordance index = 0.6) and the Ria Formosa (161 events; global P < 0.0001; ΔAICc relative to null model = -139.45; concordance index = 0.76) of success ratio in burrowing the head (a modified hazard ratio) in function of treatments and covariates when they have evidence of an effect: source temperature and target testing temperature (in °C). The time-to-burrowing is represented with boxplots for the Humber (E) and Ria Formosa (F) *per* tested individual (open symbols) and *per* treatment (closed symbols representing mean values ± 95 % confidence intervals). Data for Humber ragworms represent the effect of treatments on the residuals of covariates and is expressed as an arbitrary unit (a. u.) with lower values indicating faster times in burrowing. Significance in (C) and (F) is indicated by pairwise Wilcoxon signed rank tests.

### Using the sensory arsenal in response to pH challenges: Transcriptomic responses

Ragworms from both populations showed many DEGs in response to future ocean conditions. For the Ria Formosa population, the **current → future pH** comparison yielded 377 DEGs and the **current → SFM** comparison 339. For the Humber population, the **current → future pH** comparison yielded 1877 DEGs and the **current → future** comparison 2328. The volcano plots for all four comparisons are provided in Supplementary Figure 3. Between the **current → future pH** conditions for Humber and Ria Formosa populations as well as **current → future** condition of Humber, only 33 (1%) genes were shared (see Supplementary Figure 4). These included two solute carriers (SLC6A5; OrthoDB:1585055at7742 and SLC5A9; OrthoDB:74094at2759) as well as PXDN (Peroxidasin; OrthoDB:274545at33208), but no common functions. While we could not identify some top DEGs due to missing BLAST+ hits, UNC22 (Twitchin, OrthoDB:5303642at2759) was the most upregulated gene in both Humber comparisons **current → future** and **current → future pH.** With regards to overall functional enrichment of genes with known orthologs, the Ria Formosa **current → future pH** comparison was characterised by DEGs involved in responses to acidic conditions: the top functions containing most DEGs were solute:sodium symporter activity, ASIC channels (specifically ASIC1, OrthoDB:3415680at2759 and ASIC5, OrthoDB:3414706at2759), and stimuli-sensing channels. In addition, corticosteroid/hormone signalling was involved (Figure 5). These functions involved both significantly up- and downregulated genes (Supplementary Figure 5 and 6). In contrast, only one gene annotated as carbonic anhydrase/carbonate dehydratase (CAH14, carbonic anhydrase 1, orthoDB 3505378at2759) was downregulated in both conditions **current → future** and **current → future pH** in the Humber population. An indirect effect of low pH exposure became apparent, as the **current → SFM** condition for the Ria Formosa ragworms was characterised by a response involving genes with function as matrix metalloproteinases, in degradation of the extracellular matrix, tRNA modification and telomere extension as well as BDNF (brain-derived neurotrophic factor) regulation of GABAα transmission. Again, a complex picture emerges as all functions contain both up- and downregulated genes (Supplementary Figure 5 and 6).

**Figure 5.**
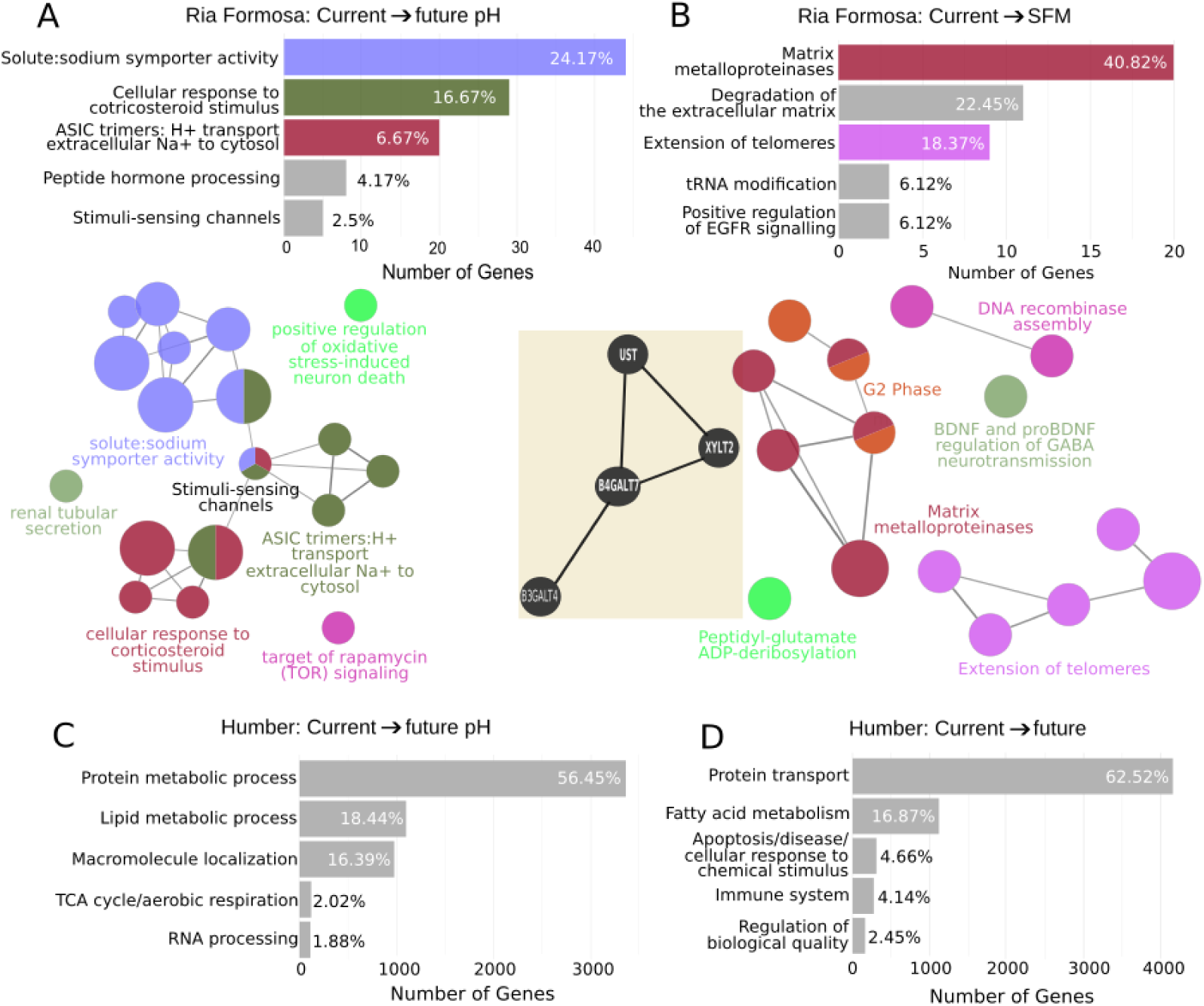
Significantly enriched functions of the annotated portion of DEGs for A) Ria Formosa ragworms responding to low pH with bar chart of most common functions by number of genes and bubble networks of connected functions; B) Ria Formosa ragworms responding to SFM with bar chart of most common functions by number of genes and bubble networks of connected functions; black inset figure in B showing a STRING network of four DEGs involved in chondroitin sulphate (“Schreckstoff”) metabolism. Bar charts of number of genes in most common functions for C) Humber ragworms responding to low pH and D) Humber ragworms responding to future conditions. Humber ragworm DEG function bubble graphs are complex owing to the large number of DEGs and thus not shown. Generated with ClueGo in Cytoscape, all terms p<0.05, analysis performed for human orthologs with the human genome as background.

In addition, STRING analysis uncovered a set of four functionally related DEGs (UST, XYLT2, B4GALT7, B3GALT4; OrthoDB IDs:5401406at2759, 4166917at2759, 306273at2759 and 532757at2759) that were involved in glycoprotein (Chondroitin / “Schreckstoff”) biosynthesis (Mathuru *et al*., 2012).

Humber ragworms showed very different gene expression patterns; besides a much larger number of DEGs, these were more functionally diverse, and lacked the expected responses to altered pH which were present in the Ria Formosa ragworms. DEGs in both comparisons were most assigned to the functions protein and lipid metabolism and transport, with the **current → future pH** treatment additionally differentially expressing genes belonging to the Citric acid cycle (TCA cycle) and RNA processing, and **current → future** treatment differentially expressing genes related to apoptosis, disease and response to chemical stimulus, immune system and homeostasis (Figure 5). Of these genes, n=105 shared the Reactome pathway HSA-1646385 “Disease”. DEG functions with large (both positive and negative) fold change in Humber **current → future pH** involve G-protein coupled receptor activity, with mostly positive log fold changes (LFCs), and response to oxidative stress (Figure 5; Supplementary Figure 5). In contrast, DEG functions with positive and negative large fold changes in Humber **current → future** treatment, respectively, differed from each other. Functions with largest positive fold changes included response to oxidative stress, immune system, responses to inorganic substances and xenobiotics, as well as G-protein coupled receptor signalling pathway. Functions with largest negative fold changes included those involved with metabolism and its diseases, regulation of hormone levels and sphingolipid metabolism (Supplementary Figures 5 and 6).

### Using the sensory arsenal in response to pH challenges: sensing and signalling

Relative to the overall expression changes within the transcriptomes, most pre-defined sensory genes were not statistically significant after FDR p-value adjustment. However, the response to low pH in both populations is characterised by absolute LFC of more than 1.5 in several genes, some of which with raw p-values <0.05 (Figure 6). Most notably, three genes annotated as SLC4A10 (Sodium-driven chloride bicarbonate exchanger orthoDB:746119at33208; abts-1 in *P. dumerilii*) had LFCs < 3 in all treatments involving low pH. In addition, OTOP was downregulated in both conditions for the Ria Formosa ragworms (orthoDB:810985at33208; a potential ortholog of Otopetrin 1), while GABR1 (Gamma-aminobutyric acid type B receptor subunit 1; orthoDB:2325at6231), GABR2, TRPA1, as well as several GPCRs, were downregulated with LFC < 1.5 in the Ria Formosa **current → future pH** condition. TRPV1, TAAR9 (trace amine-associated receptor 1-like orthoDB:2901788at2759), OPSR (green-sensitive opsin orthoDB:5350930at2759), and GBRA6 (orthoDB:4265336at2759) were upregulated with LFC > 1.5 in the Ria Formosa **current → future pH** condition. Both Humber and Ria Formosa **current → future pH** conditions differentially expressed TRPA1, which was upregulated in the former and downregulated in the latter population.

The response of Ria Formosa ragworms to chemical cues from low pH-stressed sea bream in the **current → SFM** condition overall induced a lower fold change profile of expression changes in this sensory gene panel. Several commonalities and differences emerged between this treatment and direct exposure to low pH treatment. Here, also SLC4A10 was differentially upregulated by **SFM** whilst it was downregulated in **current → future pH**, in which TRPC6 was also upregulated. Of the other pH responsive genes, the downregulation of OTOG (Otogelin, OrthoDB:1326702at33208) and OTOP is a common theme in both populations among all pH drop treatments, and in the SFM condition, likewise two copies of OTOP are downregulated. GBRA6 was upregulated in both **current → future pH** and **current → SFM** conditions. Instead, in the SFM condition, ABCB6 (ATP binding cassette subfamily B member 6; orthoDB:2876209at2759) showed downregulation. Among the GPCRs, a gene annotated as GPR45 was the most downregulated, and a range of other GPCRs including one annotated as FMAR were also downregulated. Despite mounting an overall much larger transcriptomic response than the Ria Formosa population, the Humber population ragworms expressed overall less sensory gene classes differently, with several being undetected. Most notably, this includes the complete absence of GABAα receptors as well as much less GPCRs. In contrast, this population in both treatments differentially expressed many more ABC transporters (downregulated, including three orthologs of ABCA1, orthoDB:6951at2759), TRP receptors, and pH regulation genes, compared to the Ria Formosa population.

**Figure 6.**
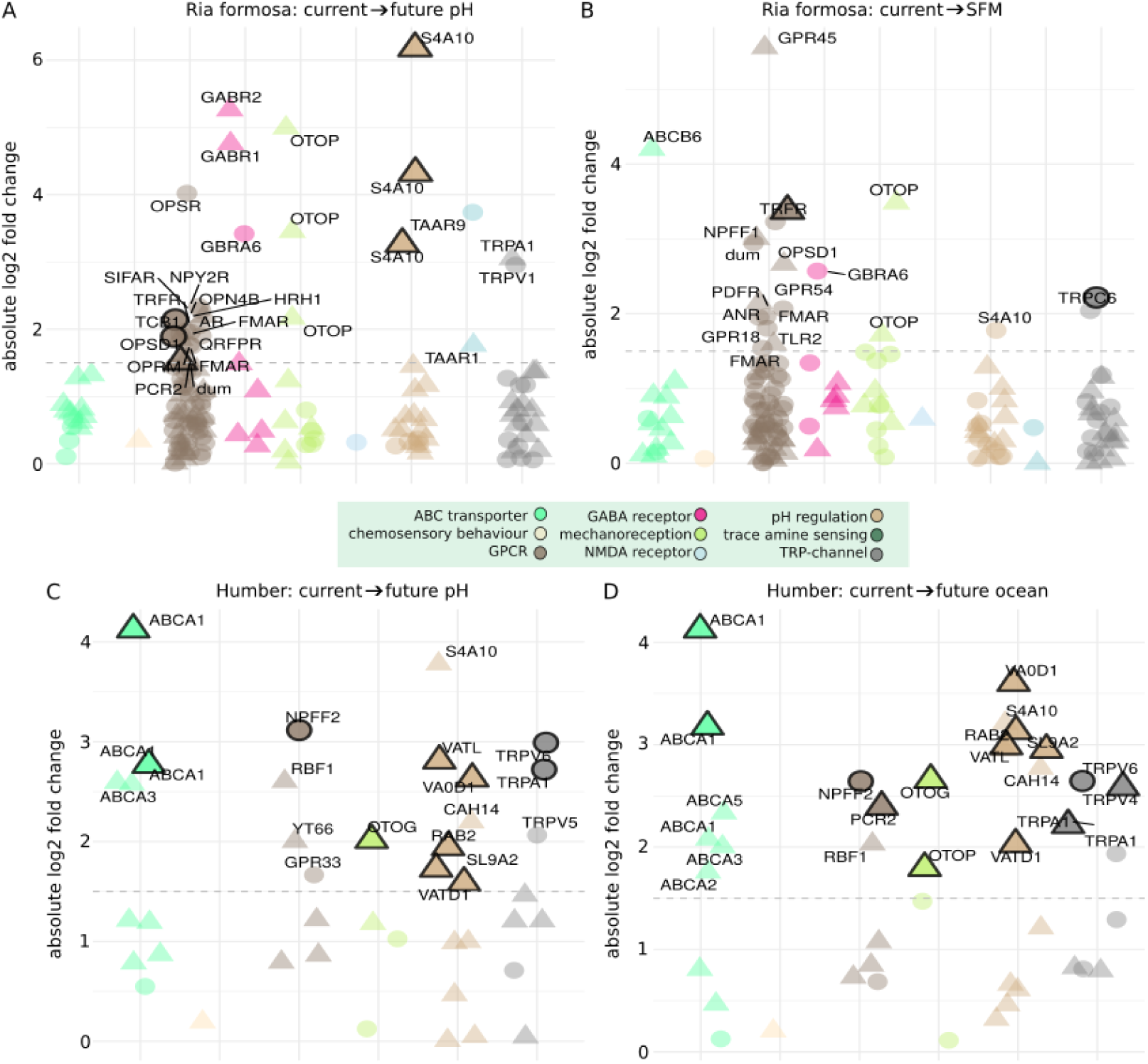
Dot plots of absolute log fold change of pre-annotated sensory function genes in response to A) future pH and B) chemical cues from low pH-exposed sea bream (current **→** SFM) in the Ria Formosa population; and C) response to future pH and D) future pH and temperature increase (current → future) in the Humber population. Genes marked with black outlines had significant raw p-values and/or absolute log2 fold-changes > 1.5 (text labels), indicating biological but not statistical significance. Triangles are downregulated DEGs while circles are upregulated DEGs.

## Discussion

### The genome of Hediste diversicolor and its molecular arsenal for pH sensing and signalling

The high cost of good quality reference genome sequencing and assembly remains a hurdle to progress in understanding the molecular responses of organisms to changes in their abiotic environment (Fernandes *et al*., 2023). Here, we provide the first annotated, chromosome resolution draft genome of *Hediste diversicolor*, an important infaunal bioturbator of coastal ecosystems, adding to this species’ molecular resources of complete mitogenome (Gomes-Dos-Santos *et al*., 2021), transcriptomes (Samico *et al*., 2022; Machado *et al*., 2024), and molecular phylogeny (Tosuji *et al*., 2019). The genome of *H. diversicolor* was most closely matched with GenBank reads from Annelida and Mollusca, since marine polychaetes (Polychaeta) are systematically placed within Annelida (Lewin, Liao and Luo, 2024). The 14 major chromosome-level scaffolds (TADs) confirm previous results from a cytological study by Leitão et al. (2010) showing a karyotype of 2n=28 chromosomes in *H. diversicolor* (Leitão *et al*., 2010). Chromosome 7 shows a minor additional TAD indicating a possibly functionally important structure which warrants further study. In addition, Chromosome 10 has lower read-to-mate pair matches with all other chromosomes, which indicates that this chromosome may be conserved, not tolerating much allelic variation, and/or be important for sex determination, in the case that the animal sequenced for the Omni-C library was of the homogametic sex. The transposable element content of 52.98% trends towards the upper range of previous % TE content assessments in twelve species of polychaetes (Filée *et al*., 2021), and is similar to the TE content of 51% in *Platynereis dumerilii* (Mutemi *et al*., 2024). Notable among the 5338 noncoding RNAs were 1596 instances of the Histone 3’UTR stem-loop which is the histone genes’ equivalent of a poly-A signal (Marzluff, 1992), and likely much higher than the actual number of histone genes in this genome, which warrants further investigation. Secondly, we only identified 25 miRNA genes which is much lower than the 580 reported for *P. dumerilii* (Mutemi *et al*., 2024) and likely because we did not have small RNA sequencing data from *H. diversicolor* to aid in miRNA identification and annotation. The gene model count of 27,839 in *H. diversicolor* after filtering corresponds to that of related polychaete species annotations of 29,000 genes in *P. dumerilii* (Mutemi *et al*., 2024), and 32,389 genes in *C. teleta* (Simakov *et al*., 2013). The SEG:MEG ratio, a measure for quality of the annotation, ranges from 2.7% in *Caenorhabdites elegans* to 18% in *Drosophila melanogaster*, and seems to be related to gene density (Sakharkar *et al*., 2004). Based on this information, the value from *H. diversicolor* falls roughly within the expected range (gene/Mb ratio of 313 against 15.6% SEGs) (Sakharkar *et al*., 2004). 14,840 genes were annotated by name against other diverse taxa, showing a high degree of gene sequence conservation. For gene model prediction, we used the transcriptomes generated from exposure to acidified pH, and cues from sea bream exposed to acidified pH, which gives us confidence that genes responding to acidified pH in this species have corresponding gene models in the genome. However, many of these genes responding to acidified pH did not have a BLAST annotation which warrants further investigation. Nonetheless, we annotated 240 instances of orthologs known for their role in sensing or signalling processes across the genome. Copy number of chemosensory gene families can be an important indicator of adaptations, such as high copy numbers of olfactory receptors for waterborne cues in habitat adaptations of green turtles *Chelonia mydas* (Bentley *et al*., 2023), fishing cats *Prionailurus viverrinus* (Bredemeyer *et al*., 2023), and across vertebrates in general (Policarpo *et al*., 2024). We found more than five copies of each OTOP, GABR1, FMAR, and GPR34 which all except GPR34 have evidence for involvement in pH sensing and signalling (see discussion and references below). Pending comparisons with more marine ragworm genomes, it is plausible to assume that the resilience of *H. diversicolor* to pH fluctuations may in part be due to gene duplications within pH-responsive pathways.

### Using the molecular arsenal in response to acidified pH

Key differences between experiments for Ria Formosa and Humber ragworms necessitated independent statistical comparisons. The Ria Formosa ragworms were field-sampled adults kept in a constant pH and temperature outdoor seawater system, while the Humber ragworms were lab-bred juveniles kept in a fluctuating pH and temperature system. RNA was sampled from whole-body Humber specimens but only from heads of Ria Formosa ragworms. Some of these experimental differences may contribute to observed differences in gene expression, but also highlight different strategies of coping with acidified pH. Humber ragworms had more DEGs, suggesting a coordinated response to conditions, though with less pronounced fold changes, indicating either a lack of acute stress response or gene expression equilibration across tissue types. Ria Formosa ragworms exhibited a typical response to low pH, involving sodium symporters and upregulation of ASIC1 and ASIC5 channels. These channels, while regulating acid-base balance, can also trigger damages to the brain and fear responses (Stelzer *et al*., 2016). Pre-defined sensory genes did not reach statistical significance in the transcriptome-wide comparison but remain biologically relevant for understanding pH sensing and -homeostasis (Dong *et al*., 2011). Subunits of the vertebrate GABAα receptor play a critical role in pH regulation of neuronal signalling (Dietrich and Morad, 2010), and are affected by environmental pH changes. CO_2_-induced behaviour changes in fish include increased anxiety (Nilsson *et al*., 2012; Hamilton, Holcombe and Tresguerres, 2014), which can be restored by blocking the GABAα receptor with GABAzine or mimicked by GABAα receptor activation through muscimol (Tresguerres and Hamilton, 2017). Similar upregulation of GABAα receptors occurs in invertebrates exposed to high CO_2_ (Moya *et ^a^l.*, 2016; Thomas *et al*., 2024). However, Ria Formosa ragworms downregulated GBRA1 and GBRA2, unlike Humber ragworms, which showed no such differential expression, suggesting adaptation to potential negative effects of GABAα receptor depolarization. This pattern aligns with findings in other species like stickleback and surfperch (Lai *et al*., 2016; Toy *et al*., 2022), species that, like the ragworms in the Humber population, also regularly experience environmental pH fluctuations. In humans and mice, loss-of-function mutations in SLC4A10 cause impaired GABAergic signal transmission, which can be rescued through the manipulation of intracellular pH (Fasham *et al*., 2023). Downregulation of this gene would conserve ATP through a reduction of the use of the Na+/K+ATPase pump (Stelzer *et al*., 2016), and further downshift GABAergic signal transmission. Pending further study, this could represent a potential adaptive mechanism to cope with sudden pH drops and associated danger of increased GABAα signalling in *H. diversicolor*, which may incur a trade-off through increased energy expenditure. The sensory gene panel in Ria Formosa ragworms revealed other upregulated potential pH-sensing candidates, such as TAAR9 (Liberles, 2015; Schirrmacher *et al*., 2021) and OPSR (Vogel and Siebert, 2001; Leung and Montell, 2017). Downregulation of the mechanoreceptor OTOP (downregulated in Ria Formosa future pH) and OTOG (downregulated in Humber future pH) might similarly be linked to intracellular pH changes (Tu *et al*., 2018). Differential expression of TRPA1 and TRPV1 in both Humber and Ria Formosa ragworms exposed to future pH (Wang, Chang and Liman, 2010; Fluck *et al*., 2022), indicates that not all responses are due to altered GABAα signalling and may involve other pathways and related consequences. Humber ragworms did not show the same acid-base balance mechanisms as the Ria Formosa population, with changes focused on metabolism, growth, and lipid/protein transport. This may be due to their juvenile stage, with acidic pH affecting development (Doo *et al*., 2012; Hue *et al*., 2022). This response mirrors that of temperate surfperch adapted to *CO*_2_ fluctuations, which upregulate metabolic activity in low pH conditions (Toy et al., 2022). Such metabolic responses to acidified seawater are common in species inhabiting pH-variable habitats, indicating that a range of species are able to mount a response to a sudden pH drop (Strader, Wong and Hofmann, 2020). Both future ocean treatments in Humber ragworms upregulated ABC transporters, particularly ABCA1, involved in lipid transport (Stelzer *et al*., 2016). Genes linked to oxidative stress were upregulated, a known response to ocean acidification (Zhang *et al*., 2012; Wong and Hofmann, 2021; Shi and Li, 2024). *Hediste diversicolor* is known to respond to acidification by increasing the antioxidant stress marker malondialdehyde (MDA), indicating heightened ROS production (Freitas et al., 2016b). The ability to mount a ROS response highlights the adaptive mechanisms that ragworms utilise in response to ROS production in acidified conditions (Sokolowski et al 2020). The Humber population may have a lower threshold for activating pH regulation channels, suggesting adaptation to variable environments according to the thermal variability hypothesis (Thompson, Crowe and Hawkins, 2002; Hofmann *et al*., 2010), and may reflect the fluctuating regime they were kept in. In contrast, GABAα signalling and GPCR expression were less altered in Humber ragworms, possibly due to whole-body RNA sampling or better innate pH adaptation.

Humber ragworms exposed to combined low pH and high temperature (future ocean conditions) exhibited a transcriptomic profile like those under the other conditions, involving general metabolism and growth, but also an oxidative stress response. However, in this population, this combined treatment also induced the expression of many genes related to apoptosis and disease. For example, TRAF2 (OrthoDB:2904475at2759), which promotes apoptosis, was upregulated, while BIRC6 (OrthoDB:117123at2759), an inhibitor of apoptosis, was downregulated (Stelzer *et al*., 2016). This suggests that future ocean conditions could promote cell fate decisions involving apoptosis. This is consistent with the observed upregulation of ATRX, PSIP1, and CHD3 (OrthoDB:2534124at33208, 2534425at33208, and 2910821at2759), which are associated with genome stability, gene expression regulation, and telomere maintenance, respectively (Stelzer *et al*., 2016). Overall, this indicates that future oceans experiencing a drop in pH in combination with increased temperature, cause serious damage to Humber ragworms, despite their pre-adaptations to pH fluctuations.

### Using the molecular arsenal in response to cues from sea bream exposed to acidified pH

Chemosensory signal transduction in polychaetes is not yet well understood (Lindsay, 2009), despite the impact of ocean acidification on these pathways becoming increasingly concerning (Roggatz *et al*., 2022). Predator detection in aquatic organisms often involves metabolic waste cues (Watson, Hamilton and Tuffnail, 2005). In polychaetes, conspecific social signals, like the sex pheromone 5-methyl-3-hep-tanone, trigger reproductive behaviours (Hardege *et al*., 1998; Hardege, 1999). Additionally, glutathione and cysteine-glutathione disulphide have been shown to induce polychaete spawning (Zeeck *et al*., 1998; Ram *et al*., 1999), and spawning can even be triggered by heterospecific cues, including those from fish (Watson *et al*., 2003).

*Nereis virens*, a closely related species to *H. diversicolor*, showed reduced activity when exposed to conspecific body extract or predator flatfish muscle extract (Watson, Hamilton and Tuffnail, 2005). In this study, water conditioned by a low pH-exposed sea bream was expected to contain a mixture of predator odours and low pH-induced metabolic cues. While these two cue types here cannot be distinguished, a concurrent study showed that exposure to this condition increased avoidance behaviours and reduced burrowing success compared to sea bream cues where fish were not pH stressed (Feugere, Angell, *et al*., 2021). Despite no direct exposure to low pH stress, several differences and similarities with the low pH treatment were observed in the stressed fish metabolite treatment. The TRPC6 (orthoDB:5406916at2759) channel, rather than TRPA1, was upregulated, suggesting shared pathways with acidified pH (Berna-Erro *et al*., 2014). Two OTOP copies were downregulated, and GBRA6 was upregulated, indicating that cues from low pH-stressed fish might activate similar signalling pathways as when directly experiencing acidified pH. The regulation of GABAα transmission was a common function of these DEGs. As a potential alternative explanation, we can rule out the presence of other cues such as social cues from conspecific ragworms present in the low pH treatment, since the system water used for all experiments was the same as that used for the control, and therefore generally present cues should not have caused any differential gene expression between control and treatments. Extracellular matrix-related genes were also prominent in this response, suggesting its role in chemical communication (Feugere *et al*., 2023). STRING analysis revealed four DEGs involved in glycoprotein biosynthesis (UST, XYLT2, B4GALT7, B3GALT4; OrthoDB IDs:5401406at2759, 4166917at2759, 306273at2759 and 532757at2759), involved in the metabolism of the skin alarm substance (“Schreckstoff”) chondroitin, which can elicit fear in fish (Mathuru *et al*., 2012). If Schreckstoff is received from the low pH-stressed sea breams, it should be tested whether it is synthesised in the ragworms, too. ABCB6 was downregulated, implicating it in the oxidative stress response (Ye, Lu and Wu, 2020). While classically associated with the mitochondrial membranes, new research also shows it to occur on the cell membrane (Rakvács *et al*., 2019). GPR45, the most downregulated gene, along with FMAR, which senses extracellular pH, suggests involvement in lipid-based chemical communication (Marchese et al., 1999). FMAR is an ionotropic receptor serving as a neuronal Na^+^ channel and related to ASIC channels which sense extracellular pH, and were upregulated in the future pH treatment (Perry *et al*., 2001; Lingueglia, Deval and Lazdunski, 2006). In the mollusc *Lymnaea stagnalis*, altering the extracellular pH has been shown to modify the sodium current of related FMARs (Perry *et al*., 2001). GPR45, related to lysophosphatidic acid (LPA) receptors, is involved in lipid signalling, potentially through the interaction of ABCB6 with membrane lipids (Marchese *et al*., 1999; Kiss *et al*., 2012; Cui *et al*., 2016; Geraldo *et al*., 2021; Lee *et al*., 2023) Lipid cues are important in chemical communication across taxa, from copepods to lizards to shore crabs (Selander *et al*., 2015; Barbero, 2016; Kamio, Yambe and Fusetani, 2022; Mangiacotti *et al*., 2023), and have been detected in cues from heat-stressed zebrafish embryos, inducing stress in receivers (Feugere *et al*., 2023).

### Burrowing response to acidified pH may reflect energetic requirements during acid-base regulation

Burrowing behaviour was fastest in both Ria Formosa and Humber ragworms when they were acclimated to and tested in acidified pH. Additionally, switching pH-acclimated ragworms to normal pH resulted in faster burrowing compared to controls, and in the Humber population, a switch from current to future pH also increased burrowing speed. In contrast, Ria Formosa ragworms acclimated to current pH and tested in acidified pH showed lower burrowing success and slower speeds (Feugere, Angell, *et al*., 2021). These results suggest that acclimation to a drop in pH has similar behavioural effects across populations, despite their different origins and experimental setups. Fast burrowing is a natural anti-predator response, reducing predation risk (Bhuiyan *et al*., 2021; Feugere, Angell, *et al*., 2021), while slow burrowing may reflect energy diverted to acid-base regulation during physiological stress, or to other stress responses like playing dead or secreting a slime cap (Feugere, Angell, *et al*., 2021). Differences in burrowing behaviour between acclimated and freshly immersed ragworms could be due to the initial cost of maintaining pH homeostasis, which decreases over time as the organism approaches a new steady state, akin to the phases of stress response (alarm stage, resistance stage, and exhaustion stage, (Chu *et al*., 2024)). The rapid burrowing response after acclimation might indicate pre-adaptation to fluctuating pH, possibly due to stable GABAα signalling. In Ria Formosa ragworms, attenuated burrowing after short-term acidified pH exposure could result from GABAα receptor downregulation and ASIC upregulation we observed. While ASIC upregulation supports acid-base regulation, it may impair neuromuscular function (Urbano *et al*., 2014). In the Humber population, rapid burrowing under low pH may be driven by the upregulation of UNC22, which regulates muscle contraction speed and force (*UniProt*, 2024)

### Conclusions

This study presents the first chromosome-level draft genome of *Hediste diversicolor*, revealing the molecular mechanisms involved in pH responses. We identified chemosensory gene families linked to pH sensing, with differing responses between populations. Our results also highlight changes in chemosensory pathways under acidified conditions, suggesting interactions between pressures and chemical communication. The Ria Formosa population showed rapid acid-base regulation, while the Humber population, composed of lab-acclimated juveniles, exhibited broader metabolic and growth changes, likely due to developmental shifts and baseline gene expression differences. These findings suggest that pH variability, environmental history and life stage strongly influence stress responses. Under future ocean conditions, the Humber population also exhibited signs of cellular stress, indicating their potential vulnerability to MHW-OAX events (Burger, Terhaar and Frölicher, 2022). Overall, this supports the idea of extending the thermal variability hypothesis to other abiotic parameters such as pH, indicating pre-adaptation to variable pH in temperate climates (Thompson, Crowe and Hawkins, 2002; Hofmann and Todgham, 2010; Hofmann *et al*., 2010; Duarte *et al*., 2013). But it also underscores the vulnerability of such populations to rising temperatures, considering the direct link between cellular processes and antipredator behaviours. While coastal organisms may be adapted to fluctuating pH, climate change will likely intensify these fluctuations (Pacella *et al*., 2018), and coinciding heat waves may be detrimental. Further research should be aimed at annotating the many genes for which we could not identify orthologs, as many of them could not be further studied in this work.

## Acknowledgements

We wish to acknowledge João Pena dos Reis, Peter Hubbard, Zelia Velez, and all staff members of the CCMAR-Ramalhete Marine Station for access to their facilities and support with the experiments. We would like to acknowledge Dr. Paola E. Campos at University College Dublin for help with phylogenetic analysis. We would like to thank Dr. Tiffany Kosch (Melbourne) for help with Synteny analysis. We acknowledge the use of pH data for the Humber Estuary kindly provided by the IMMERSE project (EU Interreg North Sea Region #945263). Cantata Bio (formerly Dovetail Genomics) performed the *de novo* assembly and HiRise scaffolding for *H. diversicolor*. JG, MS, JDH, KWV, TS, RP, SB, and CG acknowledge funding by Natural Environment Research Council (NERC grant #NE/T001577/1), and KWV, JH, and LF acknowledge funding by the University of Hull toward the MolStressH2O doctoral research cluster. KWV, KB, MP, JL, and SV acknowledge funding by the European Union (ERC, MolStressH2O, #101044202). Views and opinions expressed are however those of the author(s) only and do not necessarily reflect those of the European Union or the European Research Council Executive Agency. Neither the European Union nor the granting authority can be held responsible for them. Furthermore, we acknowledge the Viper High Performance Computing facility of the University of Hull and its support team, especially Darren Bird and Chris Collins, and the Sonic High Performance Computing facility of the University College Dublin and its support team. Lastly, we would like to thank all members of OdysysLab (Dublin and Hull) and the ChemEcolHull group for helpful discussions.

## Supplementary Materials

### Supplementary Methods

#### Genome sequencing using PacBio and proximity ligation methods

with the following workflow: Quantification of DNA samples was performed using a Qubit 2.0 Fluorometer (Life Technologies, Carlsbad, CA, USA). The PacBio SMRTbell library, approximately 20 kb in size, was constructed for PacBio Sequel sequencing using the SMRTbell Express Template Prep Kit 2.0 (PacBio, Menlo Park, CA, USA), following the manufacturer’s recommended protocol. The library was then bound to polymerase using the Sequel II Binding Kit 2.0 (PacBio) and loaded onto a PacBio Sequel II. Sequencing took place on PacBio Sequel II 8M SMRT cells. For each Dovetail Omni-C library, formaldehyde was used to fix chromatin in the nucleus. The fixed chromatin was digested with DNase I and then extracted. Afterwards, chromatin ends were repaired and ligated to a biotinylated bridge adapter, followed by proximity ligation of the adapter-containing ends. After proximity ligation, crosslinks were reversed and the DNA purified. Purified DNA was treated to remove biotin that was not internal to ligated fragments. Sequencing libraries were created using NEBNext Ultra enzymes and Illumina-compatible adapters. Biotin-containing fragments were isolated with streptavidin beads before the PCR enrichment of each library. Finally, the library was sequenced on an Illumina HiSeqX platform, aiming for ∼30x sequence coverage.

#### Genome assembly, gene prediction and genome annotation

*Wtdbg2* (Ruan and Li, 2020) was run to construct the PacBio *de novo* assembly. *Blobtools v1.1.1* (Laetsch and Blaxter, 2017) was used to identify potential contamination in the assembly based on BLAST (v2.9) (Altschul *et al*., 1990) results of the assembly against the NCBI nucleotides database. A fraction of scaffolds were identified as contaminants and were removed from the assembly. The filtered assembly was then used as an input to *purge_dups v1.1.2* (Guan *et al*., 2020) and potential haplotypic duplications were removed from the assembly, resulting in the final assembly. The input PacBio *de novo* assembly and Dovetail OmniC library reads were used as input data for *HiRise*, a software pipeline designed specifically for using proximity ligation data to scaffold genome assemblies (Putnam *et al*., 2016). Dovetail OmniC library sequences were aligned to the draft input assembly using *bwa* (https://github.com/lh3/bwa).

The separations of Dovetail OmniC read pairs mapped within draft scaffolds were analysed by *HiRise* to produce a likelihood model for genomic distance between read pairs, and the model was used to identify and break putative misjoins, to score prospective joins, and make joins above a threshold. TADs were identified using the *Arrowhead* program implemented in the *Juicertools* package (*JuicerTools*, no date). TADs were called at 3 different resolutions: 10 kbp, 25 kbp, and 50 kbp. The parameters used were -k KR -m 2000 -r 10000, -k KR -m 2000 -r 25000, and -k KR -m 2000 -r 50000. The completed assembly was submitted to NCBI and will be made available as (BioProject: SAMN43593757). The final genome assembly was then masked for repeats with *Repeatmodeler* and *Repeatmasker* (Smit, 2004; Flynn *et al*., 2020) for the purpose of gene prediction and -annotation.

For annotation, we ran the BRAKER2 pipeline for gene prediction (Brůna *et al*., 2021). First, a concatenated set of nine transcriptomes obtained from the Ria Formosa *H. diversicolor* population (see methods in RNAseq paragraph below for a description of experimental conditions control, future pH, and cues of low pH-exposed gilthead sea bream *Sparus aurata*) was scanned for contamination using *blobtools* (Laetsch and Blaxter, 2017) (Supplementary Figure 8) and then aligned to the genome using STAR (Dobin *et al*., 2013).

BRAKER2 was run first on protein evidence from Polychaeta in UniProt - OrthoDB (Kuznetsov *et al*., 2023; UniProt Consortium, 2023) and subsequently with the RNAseq alignment of *H. diversicolor*. The output files from the two runs were combined with TSEBRA (Gabriel *et al*., 2021), and BLASTp searches against the NCBI protein database were performed for initial annotation. We noticed that TSEBRA did not perform very well as several gene models with high prediction scores as well as both RNA and protein BRAKER2 evidence were not carried through to the result of TSEBRA. We therefore opted to additionally manually process the results from the RNAseq and protein BRAKER2 output files using the R packages *dplyr* (Wickham *et al*., 2014), *GenomicRanges* (Lawrence *et al*., 2013), and *rtracklayer* (Lawrence, Gentleman and Carey, 2009). First, output files were merged, and identical rows based on scaffold number, start, end, width, strand, and type, were collapsed with the rest of the data carried over from RNA evidence as priority, as that was obtained from the same species, and protein evidence as second priority. Then, the *findOverlaps* function from *GenomicRanges* was used to identify and collapse further overlapping regions between RNA and PROT gene predictions. If such overlaps were found, the higher of the two BRAKER2 scores was retained, and the evidence type column updated with RNA_PROT to denote combined evidence. Then, the protein BLAST results were added to the combined dataframe. This dataframe was merged with the output from TSEBRA based on the gene ID column. Subsequently, the scaffolds were organised by length and renamed into the 1-14 chromosomes predicted for *H. diversicolor* (see TADs plot, Figure 1). All annotations not belonging to these 14 scaffolds were removed from the annotation file (GTF) as they were both substantially smaller in size and had substantially fewer gene models. Due to merging in the annotations from the TSEBRA run, additional duplications were expected. Therefore, the removal of duplicates by identity and *findOverlaps* merge by ranges were repeated. Due to merging of RNA and PROT evidence, which resulted from independent BRAKER runs, these output files had non-matching transcript and gene IDs assigned. Therefore, BRAKER gene models were sorted by position on the chromosomes, neighbouring inconsistent gene names were unified, and gene IDs were additionally merged in from TSEBRA into a new column. That way, gene ID was in one column with ID tags originating from different lines of evidence having been unified in name and were then subsequently renumbered. Eukaryotic genes typically only have a small fraction of genes being single-exon genes, and therefore many gene models with only one CDS are likely false positives. Single exon genes <250 base pairs in length were filtered out of the GTF. In addition, BRAKER2 scores the match between RNA or protein evidence and the gene model. To be conservative, gene models with a score of <0.99 based on RNA evidence were likewise filtered out, retaining all protein BLAST hits. 1284 further gene models were removed from the GTF due to not containing any CDS prediction, which could not have been noncoding RNA genes since the RNAseq was performed on enriched mRNA.

Noncoding RNAs in the genome were identified *de novo* using *Infernal* against the Rfam database (Nawrocki and Eddy, 2013; Kalvari *et al*., 2021). This was done by using the annotated genome from BRAKER2, using easel to determine the size of the total database for the genome that was annotated (total # residues: 951207279), then using Rfam to search for the non-coding RNAs within the genome. The single-to multi-exon gene ratio (SEG:MEG ratio, also referred to as “mono-exonic/multi-exonic gene counts” or mono:multi ratios (Vuruputoor *et al*., 2023) can serve as a quality check of the annotation and was determined across all protein-coding genes.

#### Phylogenetic analysis

For phylogenetic analysis, flat priors (i.e. uniform distributions) for the substitution rate (10-12 - 12-2 substitutions/site/year) were applied. We considered a GTR substitution model with a Γ distribution and invariant sites (GTR + G + I, as best-fit model selected by BIC (Bayesian Information Criterion) with jModeltest v2.1.10 (Posada, 2008), an uncorrelated relaxed log-normal clock to account for variations between species, and a tree prior for demography of coalescent extended Bayesian skyline. The Bayesian topology was conjointly estimated with all other parameters during the Markov chain Monte-Carlo (MCMC) and no prior information from the tree was incorporated in BEAST. The analysis was run for 10 million steps and sampled every 10,000 steps, discarding the first 1,000 steps as burn-in. Convergence to the stationary, sufficient sampling (effective sample size > 200) were checked by inspecting posterior samples with Tracer v1.7.1 (Rambaut *et al*., 2018). Maximum clade credibility method in TreeAnnotator (Helfrich *et al*., 2018) was used to determine the best-supported tree. FigTree v1.4.4 (Rambaut, no date) was used for visualisation of the phylogenetic tree.

#### Transcriptomics of H. diversicolor from the Ria Formosa estuary in response to lowpH

Centrifuge tubes were placed in a polystyrene container and kept cool using ice packs to prevent heating. Rragworms were exposed for their three experimental conditions in a sequential order to allow individual handling, with the exact timing of exposure being 65 ± 5 min. At the end of the experiment, ragworms were rinsed in control seawater to remove sand. Ragworms were individually sampled in microcentrifuge tubes containing 1 mL of RNAlater^®^ and immediately frozen at -20 ℃. Ragworm specimens were shipped at room temperature in 800 µL RNAlater^®^ to the University of Hull, UK, where they were stored at -80 ℃ before further processing for downstream molecular analysis.

Ragworms from the Ria Formosa were large, compared to those from the Humber. RNA was extracted from ∼25 mg of head tissues using the RNA extraction methods modified from (Feugere *et al*., 2023). Ragworm samples were thawed, blotted on absorbent paper, and a 25 mg portion of the head was weighed and placed in a microcentrifuge tube with 1 mL ice-cold TRIzol. Ragworm samples were randomised for extraction blind to the experimenter. Ragworm tissues were homogenised for 15 s in ice using an Ultra-Turrax electric homogeniser and left for 5 min at room temperature (RT) and overnight at -80 ℃ to improve cell lysis. Upon thawing, ragworm tissues were further homogenised using a plastic pestle and via repeated passage through a p200 tip. Following 10 min of centrifuge at 12,000 g at 4 ℃, the supernatant was transferred to a new nuclease-free microcentrifuge tube. The nucleoprotein complex was let to dissociate by incubating samples at room temperature for 5 min. Next, 0.2 mL of ice-cold chloroform were added and samples thoroughly mixed by inversion, shaking, and vortexing, before another incubation at RT for 15 min. Samples were centrifuged 15 min at 12,000 rpm at RT and the aqueous phase was transferred to a new nuclease-free microcentrifuge tube. Then, 0.25 mL of isopropanol (99.5% #184130010, Acros Organics) were added and samples mixed by inversion before 10 min incubation at RT. RNA samples were incubated 10 min at room temperature and centrifuged 10 min at 12,000 rpm at RT, and the supernatant removed to obtain a pellet. The pellet was washed by resuspending it, by vortexing, in 0.5 mL ice-cold 75 % ethanol and centrifuging at 10,000 rpm at 4 ℃ for 5 min before removing all excess ethanol supernatant. The washing process was repeated two other times, and the pellet was air-dried at RT for 5 min and heated at 55 ℃ for 2 min to evaporate all excess liquid. The RNA pellet was resuspended by heating at 55 ℃ for 5 min and regular vortexing in 70 µL molecular-grade water. Contaminating DNA was removed through a DNAse I treatment (#10792877, Invitrogen™ TURBO DNA-free™ Kit) by adding 0.1 (7 µL) volume of 10X Turbo DNAse buffer and 1 µL Turbo DNAse enzyme and incubating 20 min at 37℃. 0.1 volume (7 µL) of DNAse inactivation reagent were added and samples incubated 5 min at RT before centrifugation at 10,000 rpm for 90 s. Next, 70 µL of supernatant containing DNAse-free RNA were transferred to a new nuclease-free microcentrifuge tube. RNA samples were purified using a sodium acetate washing protocol inspired from (Walker and Lorsch, 2013) to remove phenol and protein contamination to recover a cleaned RNA sample of 75 µL as described by Feugere and colleagues (Feugere *et al*., 2023). 4 µL of RNA samples were used to determine RNA concentrations using the Qubit (Qubit Broad Range kit, Invitrogen, and QubitTM fluorometer 3.0), whilst 2 µL were used to determine quality ratios using the NanoDrop 1000, and 5 µL were loaded on a 1.2 % agarose gel to check RNA integrity.

#### Whole-body Transcriptomics in H. diversicolor from the Humber

Samples were individually thawed and blotted dry to ensure no carry over of RNALater during RNA extractions. Total RNA was extracted using the crude TRIzol method following the manufacturer’s recommended protocol (BioSciences Ltd, Dublin, Ireland). Whole organisms were placed in individual sterile nuclease-free 1.5 ml Eppendorf microcentrifuge tubes (#S1615-5510, Cruinn Diagnostics Ltd, Dublin, Ireland) along with 100 µL Trizol reagent (#15596026, BioSciences Ltd, Dublin, Ireland). Tissue was then homogenised using a pellet pestle (#13236679, Fisher Scientific™, Dublin, Ireland) for 30 sec. A further 400 µL Trizol reagent was added to the tube, with a further 15-20 sec of homogenisation taking place. Samples were centrifuged for 10 min at 12,000 x g at 10°C to discard any fat and cell debris. The supernatant of each sample was transferred to a new 1.5 ml Eppendorf microcentrifuge tube before being incubated at room temperature for 5 min to permit complete dissociation of nucleoprotein complex. To the supernatant, 200 µL of ice-cold chloroform (chloroform:isoamyl, 24:1) (#C0549-1QT, Merck, Darmstadt, Germany) was added before vortexing the mixture. Samples were then incubated at room temperature for 15 min before centrifugation for 15 min at 12,000 x g at room temperature. The aqueous (upper) phase was then transferred to a new 1.5 ml Eppendorf microcentrifuge tube, ensuring the other phases (lower red phenol-chloroform and interphase) were not disturbed. To each sample, 250 µL of room temperature isopropanol (#10215390, Fisher Scientific, Dublin, Ireland) was added before being inverted several times to mix. Samples were then incubated for 10 min at room temperature before being centrifuged at 12,000 x g for 10 min at room temperature. The supernatant was then removed from each tube before the washing of each sample’s pellet could take place. Each pellet was then resuspended in 500 µL ice-cold 75% Ethanol (#12438760, Fisher Scientific™, Dublin, Ireland) before being vortexed briefly and then centrifuged at 10,000 x g for 5 min at 4°C. After centrifugation, the supernatant (ethanol) was discarded, and the washing with 75% ethanol was repeated two more times ensuring all excess ethanol was removed at the end of the washing phase. Samples were then left to air dry at room temperature for 3 min before being heated at 55°C for 2 min, making sure all the ethanol had evaporated. Each sample’s pellet was then resuspended in 50 µL of molecular-grade water (#10307052, Fisher Scientific™, Dublin, Ireland) before being heated at 55°C for 5 min and finally being vortexed thoroughly. Samples were then immediately placed on ice.

Carryover DNA was removed using a Invitrogen™ TURBO DNA-free^TM^ Kit (#AM1907, BioSciences, Dublin, Ireland). For this, 0.1 volume of 10X Turbo DNAse buffer along with 1 µL DNAse enzyme were added to each RNA sample, gently mixed, and allowed to incubate at 37°C for 20 min. Resuspended DNAse inactivation (0.1 volume) was added to each sample before being mixed thoroughly and incubated for 5 min at room temperature (occasionally mixing each sample by flicking the Eppendorf microcentrifuge tube). Samples were then centrifuged at 10,000 x g for 90 sec. A known volume of RNA sample was then transferred to a new 1.5 ml Eppendorf microcentrifuge tube before conducting the sodium acetate protocol modified from (Walker and Lorsch, 2013) for the removal of phenol carryover. 3M sodium acetate was prepared by adding 24.6 g sodium acetate anhydrous (#11347267, Fisher Scientific™, Dublin, Ireland) in 80 ml molecular grade water before being adjusted to a pH of 5.2 with glacial acetic acid (#11475160, Fisher Scientific™, Dublin, Ireland) and completing the volume to 100 ml with molecular grade water. Next, 0.1 volume of 3M sodium acetate was added to each sample and thoroughly mixed, before 2.5 volumes of 100% ethanol were added. Samples were then incubated at -20°C for 24hrs. After incubation, the RNA was pelleted by centrifugation at 12,000 x g for 15 min at 4°C. supernatant was then carefully removed from each sample, and the RNA pellet washed by adding 2.5 volumes of 75% ethanol and allowing the pellet to soak for 2 min before centrifugation at 12,000 x g for 2 min at 4°C and removing all ethanol. This washing process was repeated a further two times before allowing the pellet to dry at room temperature (or heated for 2 min at 55°C). RNA pellets were then dissolved in molecular grade water by vortexing thoroughly.

Aliquots of 10 µL were taken for quality and quantity assessment, with the remaining 40 µL being stored at -80°C prior to being sent in dry-ice to NERC Environmental Omics Facility for RNA-sequencing. RNA purity was assessed using the BioDrop µLite Spectrophotometer (BioDrop, Cambridge, England). All RNA samples contained a volume of 50 µL with > 2,700 ng RNA, and acceptable purity shown by the 260/280 and 260/230 ratios respectively >1.9 and > 1.9. Sequencing was done by Liverpool Genomics (UK) involving preparation of dual-indexed, strand-specific RNASeq library using the NEBNext polyA selection and Ultra II Directional RNA library preparation kits, followed by Illumina NovaSeq sequencing (paired-end, 2x150 bp).

## Supplementary Figures

**Supplementary Data Sheet:** <u>XXX will be made available with publication XXX</u>

**Supplementary Figure 1.**
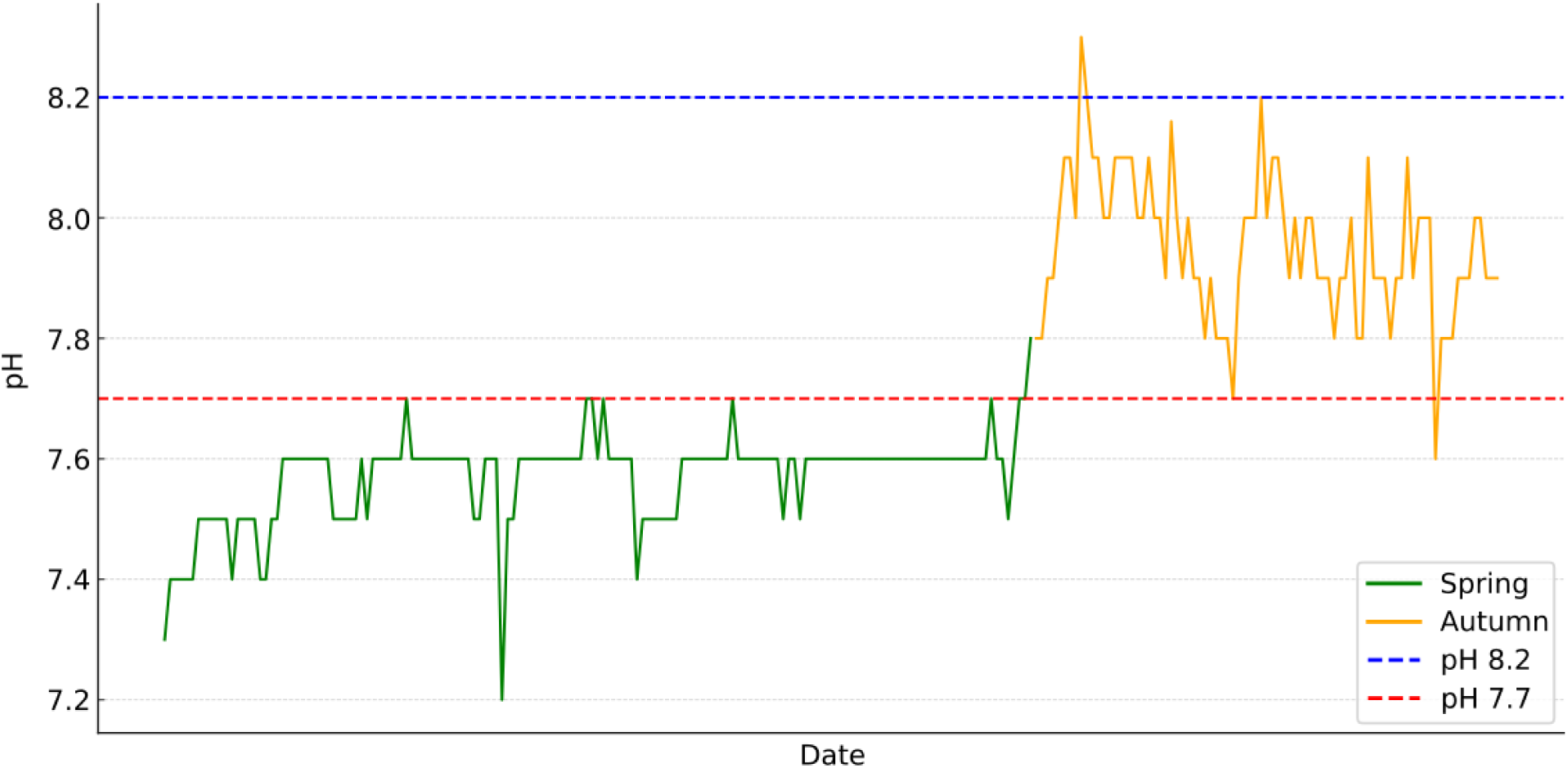
Natural pH fluctuation of the Humber estuary near Grimsby in Spring and Autumn 2021, at the time when ragworms were collected however more downstream. The average pH was 7.57 in Spring and 7.95 in Autumn.

**Supplementary Figure 2.**
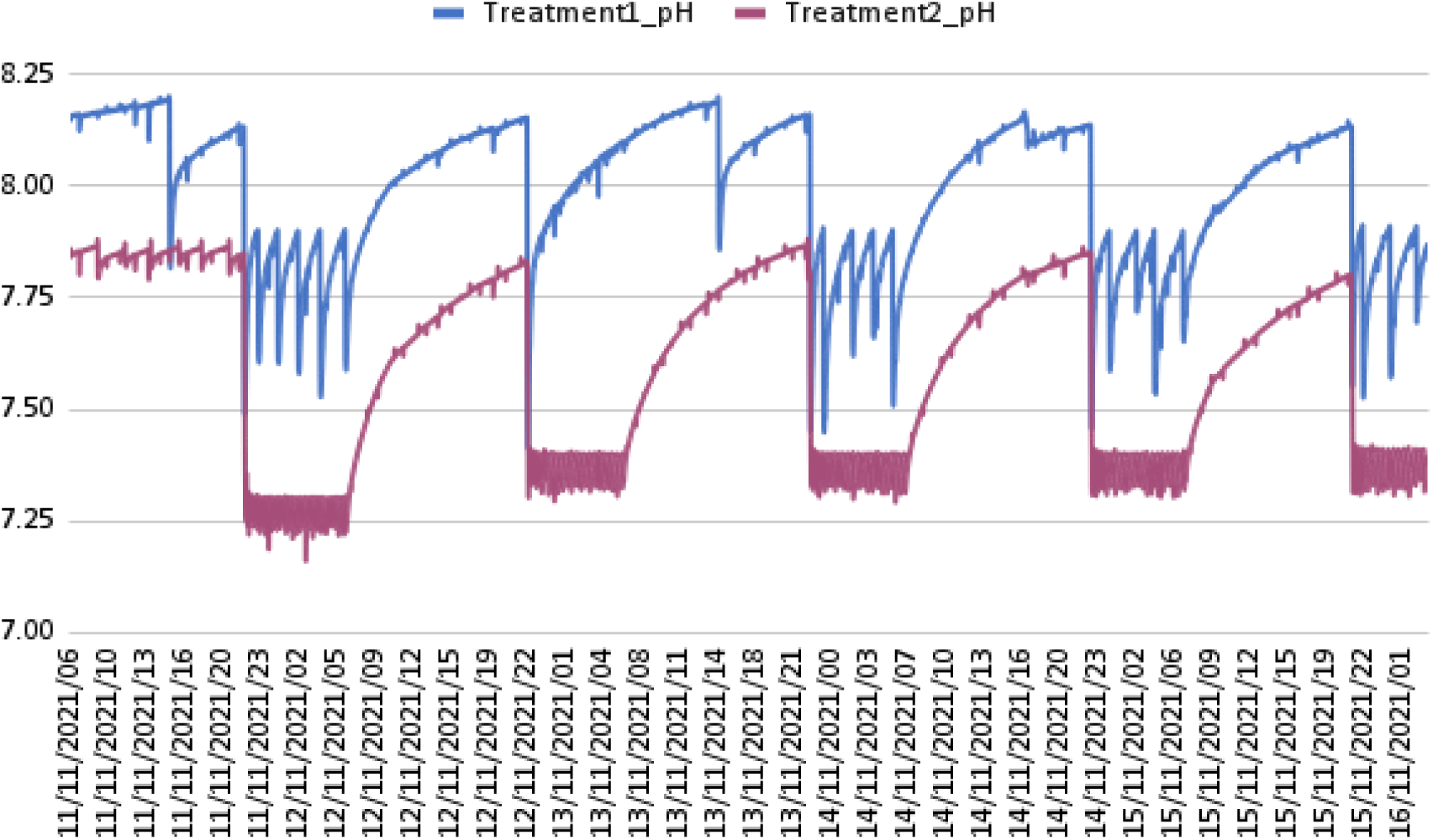
Diel pH fluctuation in long-term treatments “current” and “future” for marine ragworms collected from the Humber estuary near Grimsby in Spring and Summer 2021.

**Supplementary Figure 3.**
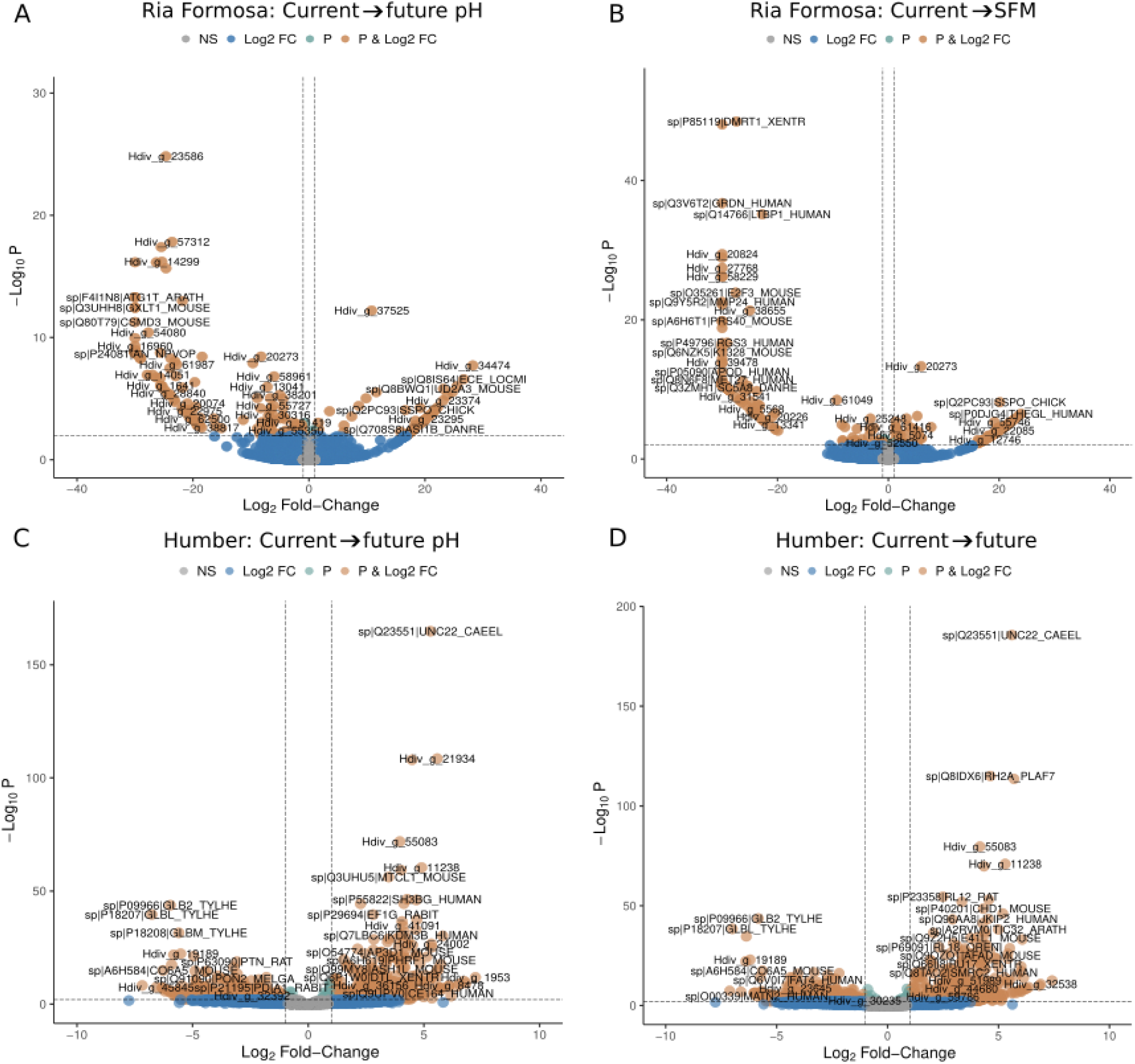
Volcano plots for gene expression differences based on 14 chromosomal scaffolds of marine ragworms faced with pH-related challenges. A) field-sampled Ria Formosa ragworms (head) switched to future pH and B) exposed to low pH-stressed sea bream cues (current → SFM). C) F2 ragworms from Humber (whole body) switched to future pH and D) to future conditions (low pH and high temperature).

**Supplementary Figure 4.**
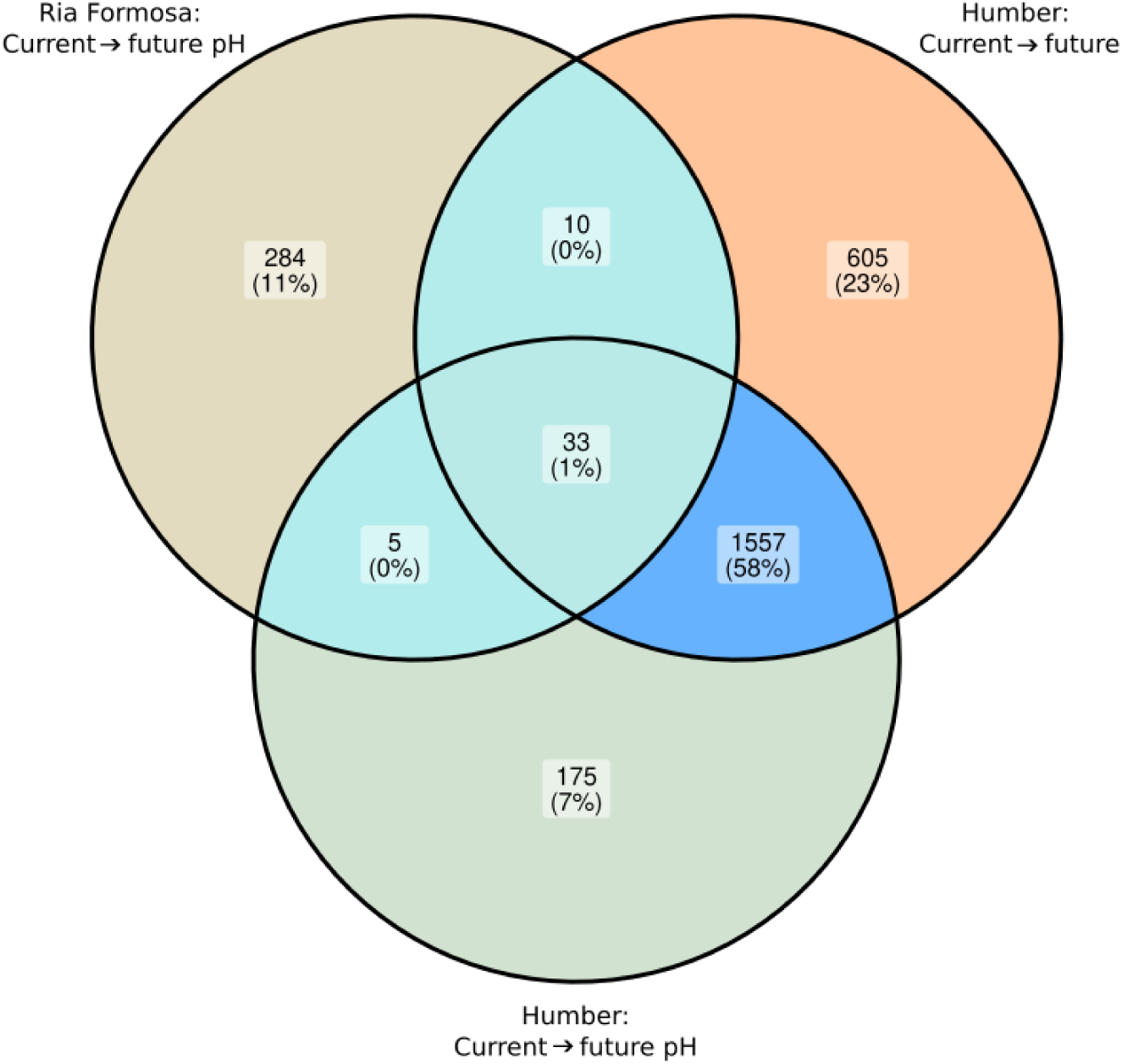
Venn diagrams to compare differential gene expression shared between conditions between Ria Formosa and Humber.

**Supplementary Figure 5.**
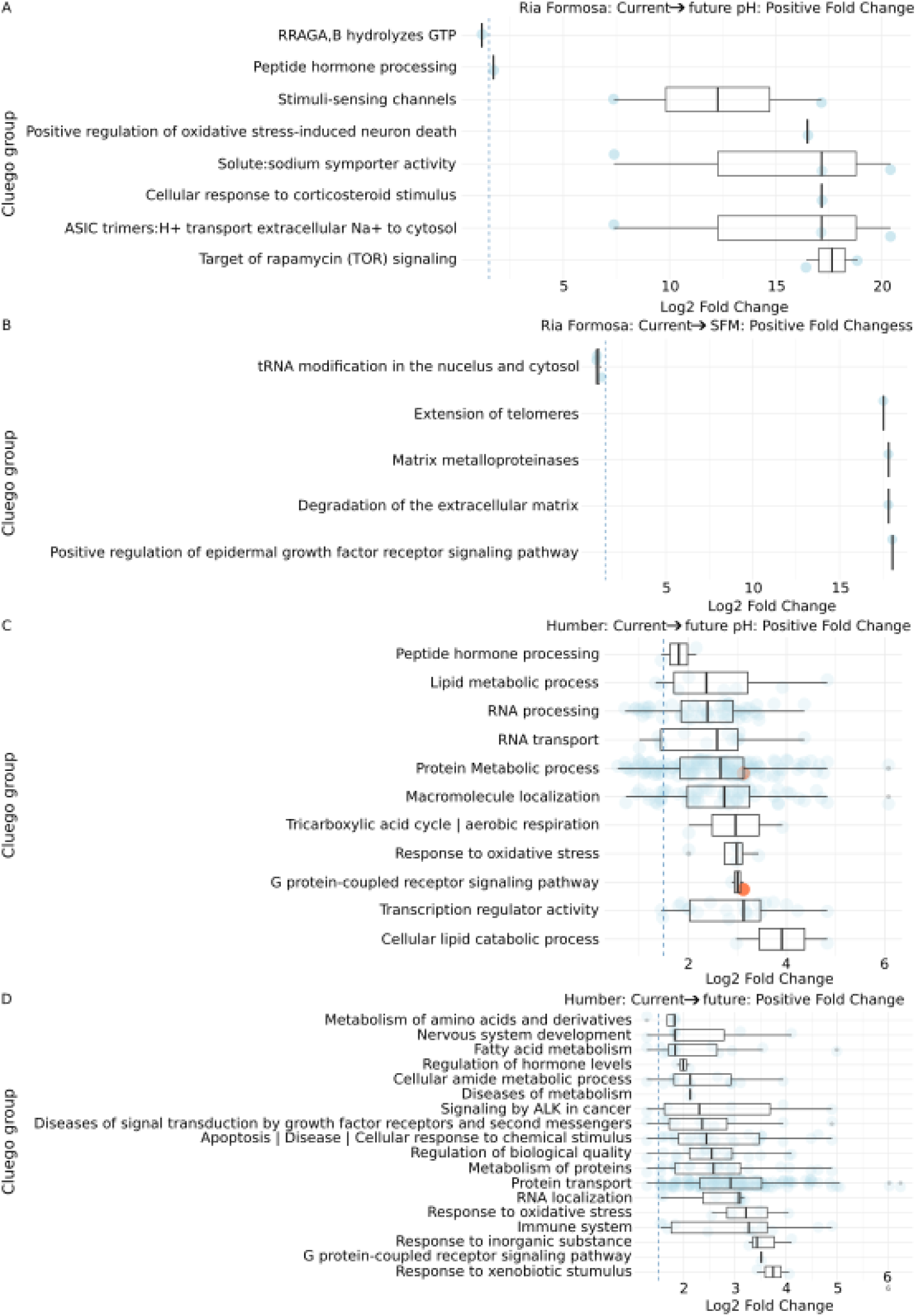
Positive fold changes of annotated significant DEGs allocated to GO term groups across the different treatments. Opposite coloured symbols represent genes with “sensory function” annotation. A) Ria Formosa, Portugal, current conditions compared to future pH. B) Ria Formosa, Portugal, current conditions compared to SFM. C) Humber, UK, current conditions compared to future pH. D) Humber, UK, current conditions compared to future.

**Supplementary Figure 6.**
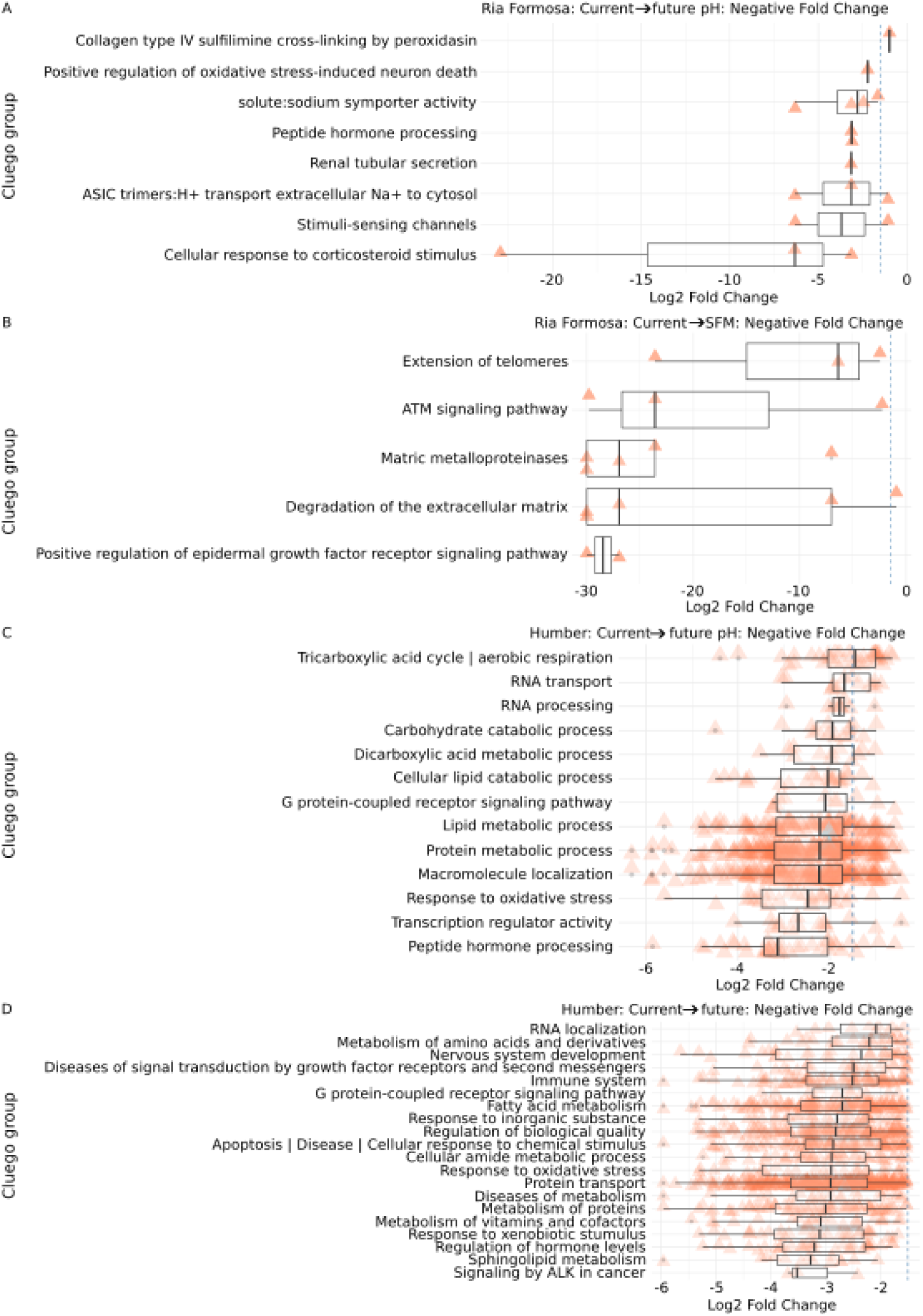
Negative fold changes of annotated significant DEGs allocated to GO term groups across the different treatments. Opposite coloured symbols represent genes with “sensory function” annotation. A) Ria Formosa, Portugal, current conditions compared to future pH. B) Ria Formosa, Portugal, current conditions compared to SFM. C) Humber, UK, current conditions compared to future pH. D) Humber, UK, current conditions compared to future.

**Supplementary Figure 7.**
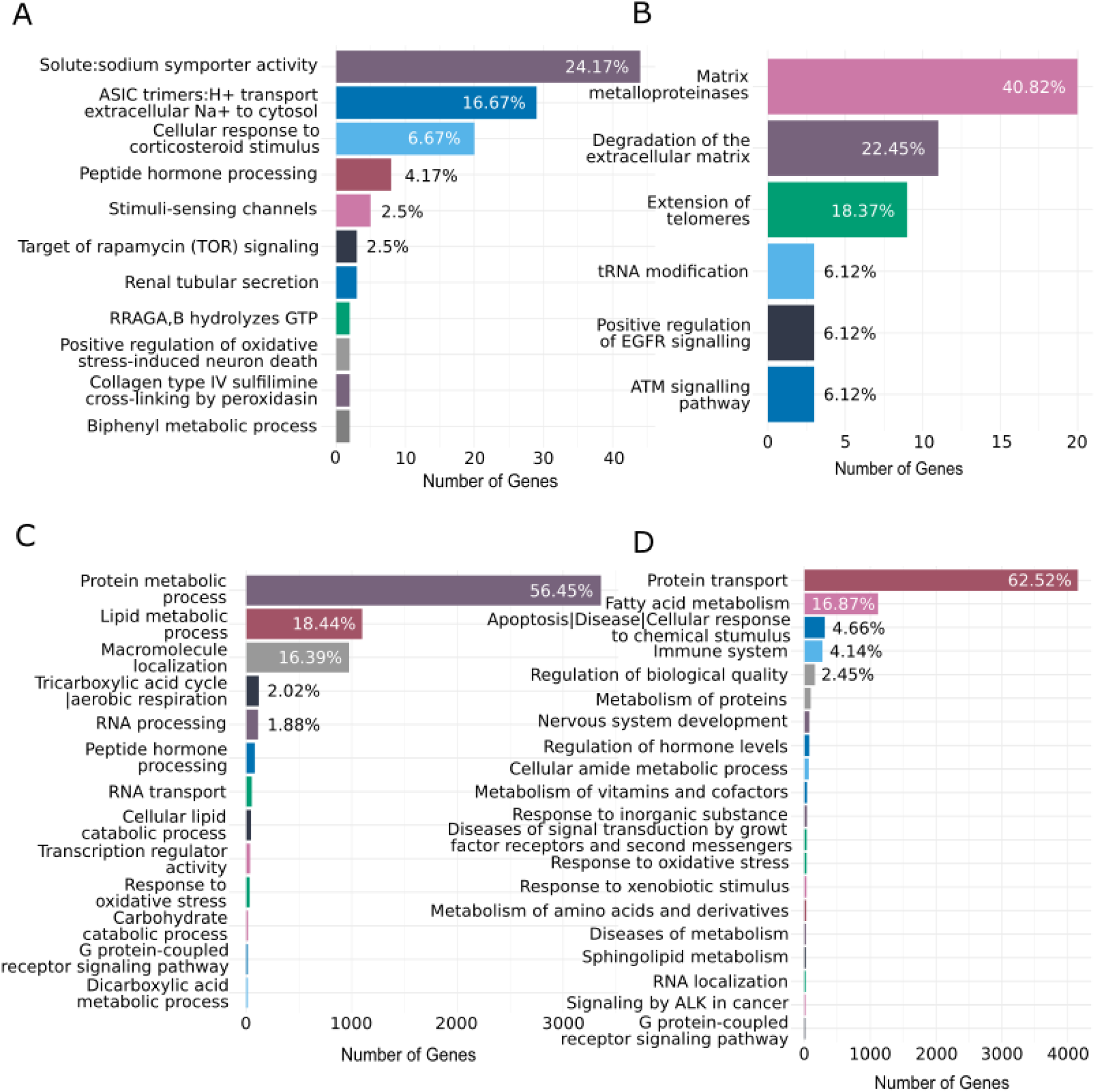
CLUEGO bar charts for all significant terms. A) Ria Formosa, current → future pH, B) Ria Formosa, current → SFM; C) Humber, current → future pH, D) Humber, current → future.

**Supplementary Figure 8.**
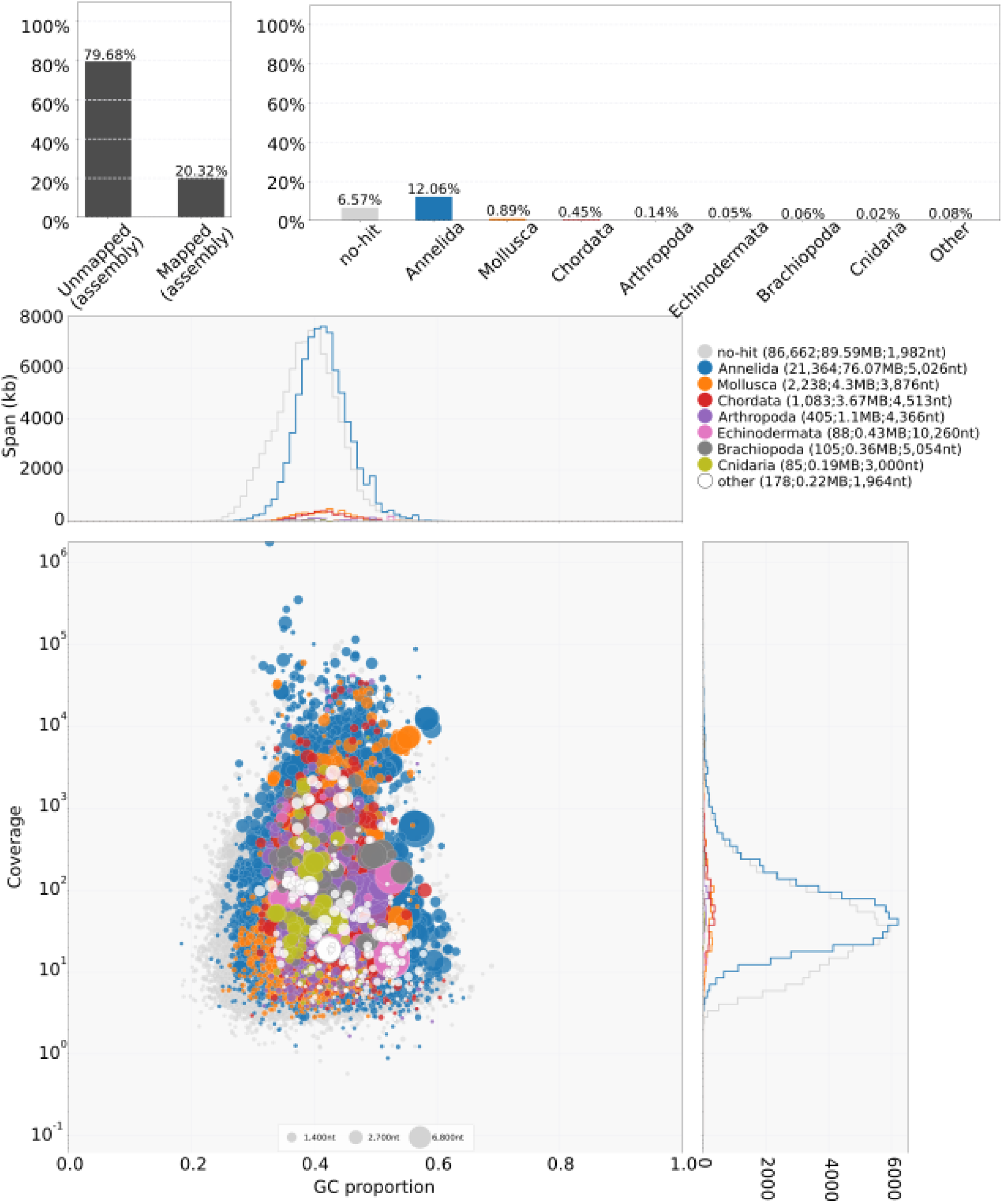
Transcriptome BlobPlot of Ria Formosa ragworms that were used for genome annotation to test for contamination with different species.

## Supplementary Tables

**Supplementary Table 1.**
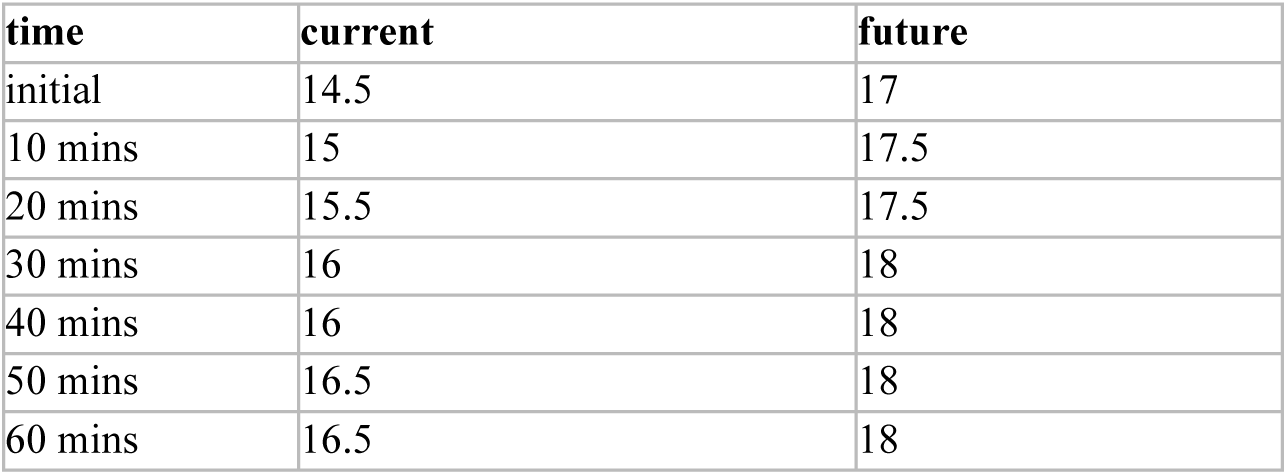
Water temperature acclimation during the behaviour experiments (within 10 mins) / sampling for small RNAseq (60 minutes).

**Supplementary Table 2.** Summary of Cox-proportional hazard ratios (HR, the exponentiated coefficient) for the burrowing success behaviour of *H. diversicolor* from the Ria Formosa (n = 189, with 161 events) or the Humber (n = 235, with 161 events). SE: standard error of coefficient, CI: confidence interval of HR. P-values indicated in bold are significant at α = 0.05. The Chi-squared (χ^2^, with its p-value) and ΔAkaike Information Criterion (ΔAICc) values are indicated relative to the null model for each population.

| Term | Coefficient | SE | Z | HR | 95% CI | P |
| --- | --- | --- | --- | --- | --- | --- |
| **Ria Formosa** ($\chi^2 = 139.4459$, $P < 0.0001$, ΔAICc = 133.32) | | | | | | |
| Treatment | - | - | - | - | - | - |
| Current (reference) | - | - | - | - | - | - |
| **Current → Future pH** | -1.0885 | 0.2964 | -3.6729 | **0.34** | 0.19-0.60 | **< 0.001** |
| **Future pH → Current** | 1.6739 | 0.2709 | 6.1797 | **5.33** | 3.14-9.07 | **< 0.001** |
| **Future pH** | 1.6387 | 5.1485 | 7.9913 | **5.15** | 3.44-7.70 | **< 0.001** |
| **Humber** ($\chi^2 = 36.5517$, $P < 0.0001$, ΔAICc = 24.18) | | | | | | |
| Treatment | - | - | - | - | - | - |
| Current (reference) | - | - | - | - | - | - |
| **Current → Future** | 1.3015 | 0.317 | 4.106 | **3.67** | 1.97-6.84 | **< 0.001** |
| **Future → Current** | 0.9765 | 0.2931 | 3.332 | **2.66** | 1.49-4.72 | **0.0009** |
| **Future** | 0.953 | 0.3387 | 2.813 | **2.59** | 1.34-5.04 | **0.0049** |
| **Worm use** | -0.4152 | 0.1091 | -3.807 | **0.66** | 0.53- 0.82 | **0.0001** |
| Source T°C | -0.1151 | 0.0744 | -1.547 | 0.89 | 0.77-1.03 | 0.1219 |
| Target T°C | -0.1918 | 0.1036 | -1.852 | 0.83 | 0.67-1.01 | 0.0640 |

**Supplementary Table 3.**
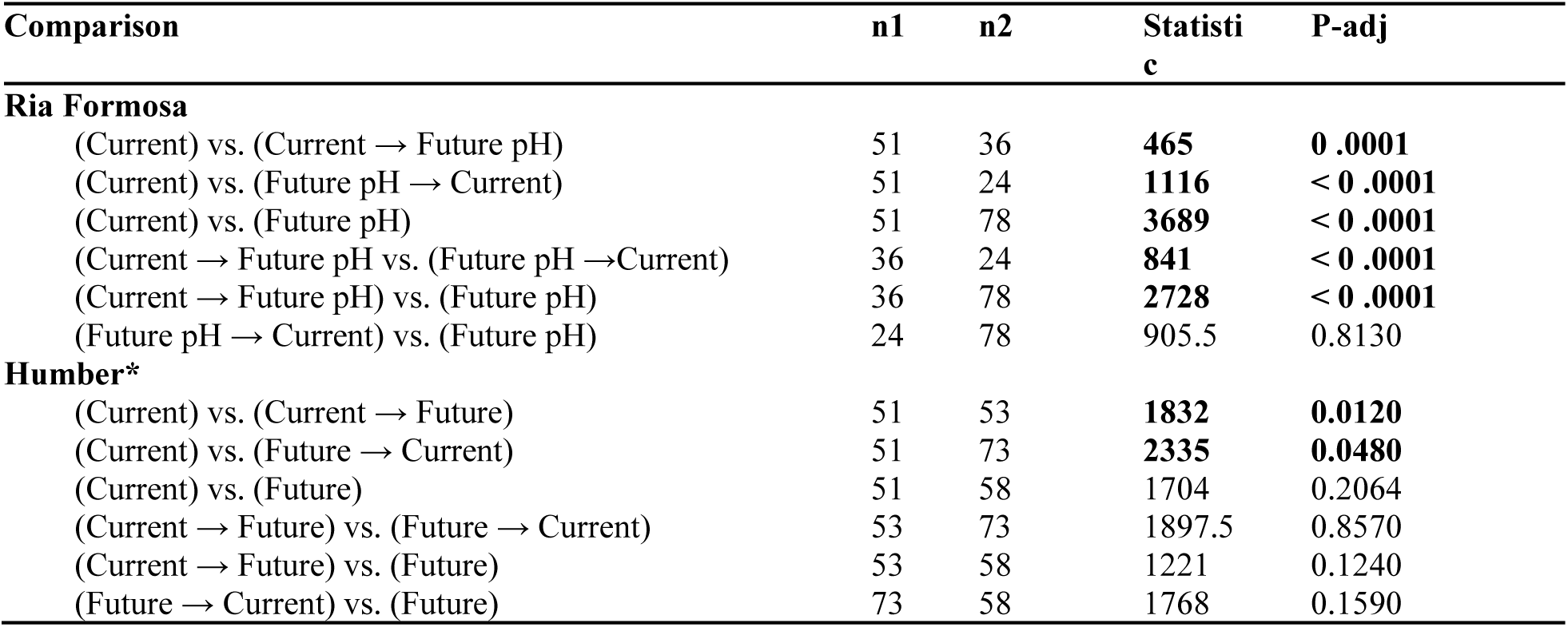
Pairwise comparisons of time taken to burrow the head in the sand in. P-values indicated in bold denote statistical significance. P-values are corrected for multiple testing using False Discovery Rate adjustment. Sample sizes are indicated for each group as (n1) vs. (n2). Statistics are from two-samples Wilcoxon tests. *Data for the Humber was analysed on the residuals corrected for the effects of covariates (worm use, source and target temperatures) to test for the effect of treatments.

